# Epigenomic analysis of hepatocellular carcinoma reveals aberrant *cis*-regulatory changes and dysregulated retrotransposons with prognostic potentials

**DOI:** 10.1101/2024.09.16.613254

**Authors:** Clooney C. Y. Cheng, Ming Fung Cheung, Ah Young Lee, Qiong Wu, Savio Ho-Chit Chow, Julie Y. J. Ang, Ignacio Riquelme Medina, Grace Lo, Haoran Wu, Weiqin Yang, Paul B. S. Lai, Kevin Y. Yip, Alfred S. Cheng, Danny Leung

## Abstract

Hepatocellular carcinoma (HCC) exhibits widespread epigenetic alterations, yet their impact on *cis*-regulatory elements (CREs) and retrotransposons remains poorly understood. Here, we present an integrated epigenomic and transcriptomic analysis of HCC tumors and matched tumor-adjacent normal tissues. We identified extensive DNA hypomethylation coupled with changes in histone modifications at partially methylated domains, CREs, and retrotransposons. These epigenetic aberrations were associated with dysregulated expression of genes involved in cell cycle regulation, immune response, and extracellular matrix organization. Notably, our findings revealed a novel mechanism for the transcriptional dysregulation of *GPC3*, a key HCC biomarker and immunotherapeutic target. We observed that *GPC3* upregulation is driven by both the reactivation of a fetal liver super enhancer and hypomethylation of *GPC3*-associated CpG islands. Moreover, we found that DNA hypomethylation-driven aberrant expression of retrotransposons carries prognostic significance in HCC. Patients with high expression of a long non-coding RNA driven by a HERVE-int element exhibited more aggressive tumors, poorer clinical outcomes, and molecular features associated with favorable immunotherapy response. Together, our study provides a comprehensive resource for understanding the role of epigenetic dysregulation in HCC and identifies retrotransposon-associated transcripts as potential biomarkers.

---

Hepatocellular carcinoma (HCC) is the major form of primary liver cancer and the third-leading cause of cancer-related deaths worldwide^1^. Alcohol abuse, chronic hepatitis B virus (HBV) and hepatitis C virus (HCV) infections, and non-alcoholic fatty liver disease (NAFLD) are key risk factors for HCC^2^. These conditions contribute to cirrhosis that often precedes HCC development^2^. While recent advances in targeted therapies and immunotherapies have improved patient disease outcome, the 5-year survival rate of HCC remains less than 20% due to frequent tumor relapse and resistance to treatments^2–5^.

Previous epigenomic and transcriptomic analyses have revealed dysregulated pathways in HCC^6–8^. These gene expression programs are modulated by non-coding sequences known as *cis*-regulatory elements (CREs), including enhancers and promoters^9^. CRE activation is coupled with epigenetic changes, including reduced DNA methylation, increased chromatin accessibility, and enrichment of specific histone modifications^10–13^. Several studies have examined the impact of epigenetic alterations on the activities of individual CREs in HCC. For example, DNA hypomethylation of an enhancer promotes C/*EBPβ* overexpression and tumor progression^14^. Moreover, in approximately 50% of HCC patients, the *CDKN2A* promoter is DNA hypermethylated, leading to its transcriptional repression and is associated with perturbed cell cycle regulation^6,15^. While changes in CRE activities have been reported in HCC, the associated epigenomic dysregulations have not been fully delineated^16–21^.

Retrotransposons are repetitive elements that constitute almost half of the human genome and provide a rich repertoire of CREs in both normal development and in diseases^22–28^. While most are epigenetically silenced, some can become derepressed in cancers and influence nearby gene expression^22,24–26,29^. Specifically, long-terminal repeat (LTR) retrotransposons, including endogenous retroviruses (ERVs), can produce transcripts and serve as CREs that have direct impacts on cancer development. In HCC, high expression of ERV-driven non-coding transcripts has been associated with HBV etiologies and less differentiated states^30^. ERV-driven long non-coding RNAs (lncRNAs) could promote DNA double-strand break repair and upregulate tumor metastasis-related pathways^31–34^. Moreover, ERV-derived CREs were reported to affect the transcriptomes of diverse cancer types^24,26^. Despite reports of retrotransposon dysregulation in HCC, its scope, molecular mechanisms, functional consequences, and clinical relevance remain largely unexplored.

Here, we deciphered the transcriptional regulatory landscape of HCC by mapping the transcriptomes, histone modification profiles, and DNA methylomes of HCC tumors and matched tumor-adjacent normal controls. Integrating the multi-modal datasets, we dissected the epigenomic mechanisms underlying aberrant activities of CREs and retrotransposons in HCC. We observed wide-spread epigenetic aberrations including global DNA hypomethylation at partially methylated domains (PMDs) and focal DNA hypomethylation at specific CREs. These changes were coupled with alterations of other epigenetic modifications. Notably, we demonstrated the interplay between super enhancer (SE) reactivation and DNA hypomethylation of CpG islands in the transcriptional regulation of glypican-3 (*GPC3*), a diagnostic marker and immunotherapeutic target for HCC^35–38^. Furthermore, we showed that high expression of a HERVE-int element-driven lncRNA in patients was associated with poorer patient survival and molecular signatures of better response to immunotherapy.

## RESULTS

### Integrative analyses reveal extensive epigenomic reprogramming in HCC

To analyze the epigenomic and transcriptomic changes in HCC, we performed stranded total RNA sequencing (RNA-seq), chromatin immunoprecipitation followed by sequencing (ChIP-seq) for four histone modifications, and whole-genome bisulfite sequencing (WGBS) on seven pairs of HCC tumors and matched tumor-adjacent normal tissues (TAN) (**Fig. 1a**). All enrolled patients are HBV positive and all except one show cirrhosis (**Supplementary Table 1**). The profiled histone modifications demarcate active CREs (histone H3 lysine 27 acetylation (H3K27ac) and histone H3 lysine 4 trimethylation (H3K4me3)) and repressed regions (histone H3 lysine 9 trimethylation (H3K9me3) and histone H3 lysine 27 trimethylation (H3K27me3))^10–12^. Importantly, we generated deeply sequenced DNA methylome maps (46X mean genome coverage and 56X mean CpG coverage, respectively, **Supplementary Table 2**). All datasets are in accordance with the International Human Epigenome Consortium (IHEC) quality standards (**Supplementary Tables 2**)^39^. To further bolster our analyses, we incorporated four publicly available DNA methylomes for TAN and imputed the absent ChIP-seq datasets, which were not generated due to insufficient sample materials^40^.

**Figure 1.**
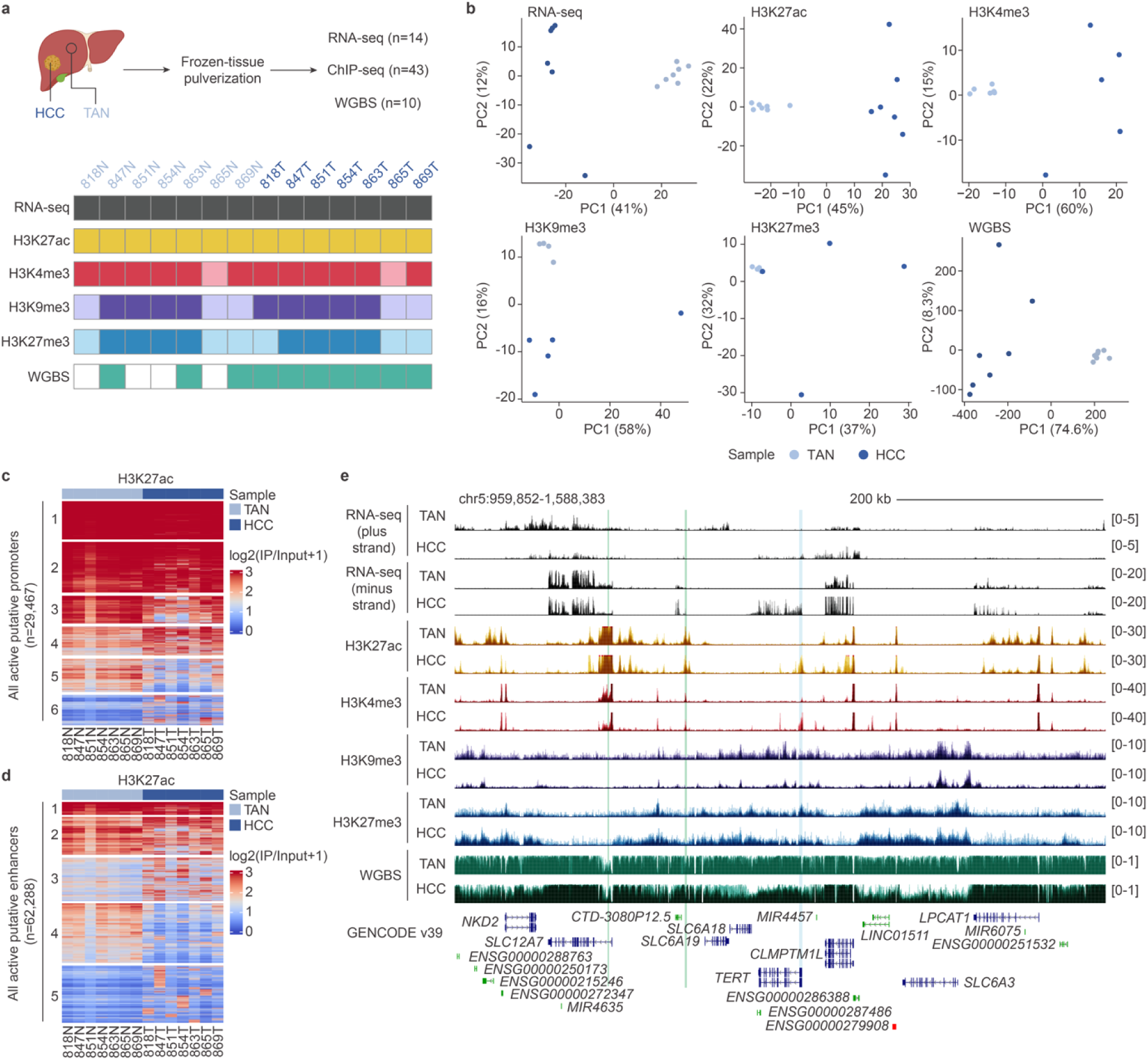
Multimodal characterization of the epigenomic landscape of HCC. **a**) Schematic of experimental design. Frozen patient-derived HCC or TAN tissue samples were pulverized and subjected to stranded total RNA-seq, ChIP-seq for the histone modifications (H3K27ac, H3K4me3, H3K9me3 and H3K27me3), and WGBS. For marks that were not profiled due to limited samples, data was imputed using Avocado. Shading of cells indicate experimental (dark) or imputed (light) datasets. **b**) Principal component analyses of the six types of profiled data. Each dot represents a sample, and the color corresponds to sample type (HCC: dark blue; TAN: light blue). **c**-**d**) Heatmap of H3K27ac ChIP-seq signals (log2-transformed fold enrichment over input) of all defined active putative enhancers (n=62,288) (**c**) and all defined active putative promoters (n=29,467) (**d**). Each row represents an element, and each column represents a sample. **e**) Genome browser screenshot showing an expanded genomic region centered at the *TERT* gene. Transcriptional upregulation of *TERT* was observed in HCC samples, coupled with the gain of active promoter epigenetic signatures (cyan) and activation of associated distal enhancers (green), which were previously defined. Stranded total RNA-seq tracks are displayed as transcripts per million (TPM), ChIP-seq tracks are displayed as fold enrichment over input, and WGBS tracks are displayed as methylated CpG to unmethylated CpG ratio (mCG/CG). All tracks are shown as composite signals by overlaying tracks across samples.

Principal component analyses (PCA) of RNA-seq, ChIP-seq and WGBS datasets show clear separation of HCC and TAN samples (**Fig. 1b**). We defined active putative promoters (n=29,467) as regions enriched with both H3K27ac and H3K4me3 peaks and active putative enhancers (n=62,288) as H3K27ac peaks that do not overlap with H3K4me3 peaks or GENCODE v39 transcription start site (TSS) definitions (±1kb flanking TSS) (**Fig. 1c**,**d**). Consistent with previous reports, we found higher variation of enhancer activities as compared to promoters both across individual samples and disease states (**Fig. 1c**,**d**)^10,13,16^. Together, we established a compendium of epigenomic and transcriptomic datasets for in-depth investigation of the regulatory landscape in HCC (**Fig. 1e**).

### Global DNA hypomethylation and focal DNA hypermethylation in HCC

Global DNA hypomethylation in intergenic regions and focal DNA hypermethylation of specific gene promoters have been reported in HCC^6,15,41^. However, these studies have largely been focused on specific loci or do not have sufficient depth and resolution^17,19^. With deeply sequenced WGBS datasets, we were able to analyze the genome-wide DNA methylation profiles of HCC at individual CpG resolutions. In agreement with previous findings, HCC exhibited significant global reduction in DNA methylation relative to TAN (**Fig. 2a**, **Extended Data Fig. 2a** and **Supplementary Table 2**)^6,15,19,41^. The loss of methylated CpGs (mCG) occurred irrespective of genomic contexts, including at CpG islands (CGIs), genic features, CREs, CTCF sites (**Fig. 2b**), and repetitive elements (**Fig. 2c**). Focal increase of mCG was also detected (**Fig. 2d**). We further defined differentially methylated regions (DMRs) in HCC, identifying 1,193 hypermethylated regions (hyperDMRs) and 128,325 hypomethylated regions (hypoDMRs) (**Supplementary Table 4**). HyperDMRs were enriched in gene promoters and genic features including UTRs and exons (**Fig. 2e**). Genes with hypermethylated promoters were associated with gene ontology (GO) terms for developmental processes (**Extended Data Fig. 2b**), which is consistent with previous observations that developmental gene promoters are frequently hypermethylated in cancers^42,43^. These genes include *CDKN2A*, a tumor-suppressor gene that blocks G1-S phase transition, amongst others^6,15,41^.

**Figure 2.**
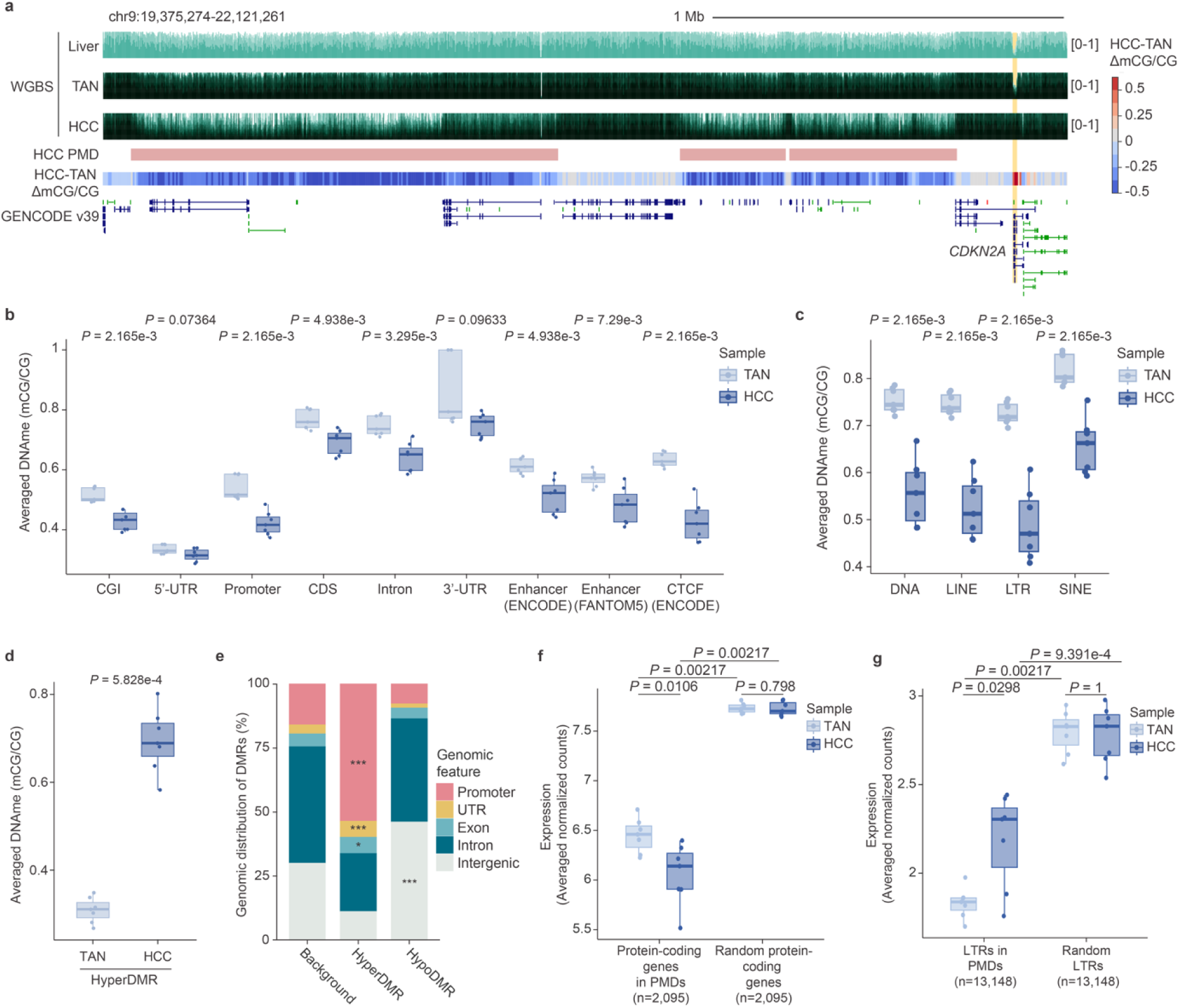
Global DNA hypomethylation and focal DNA hypermethylation in HCC. **a**) Genome browser screenshot showing an expanded genomic region surrounding the *CDKN2A* gene. This illustrates the phenomenon of global DNA hypomethylation and focal DNA hypermethylation in HCC. PMDs defined in HCC are annotated by pink bars (HCC PMD). The averaged DNA methylation difference between HCC and TAN per 5-kb window is shown as heatmap track (HCC-TAN ΔmCG/CG). *CDKN2A* promoter is highlighted by yellow shading. WGBS tracks are displayed as mCG/CG. Liver WGBS dataset from Roadmap Epigenomics Project is shown as single replicate. Other tracks are shown as composite signals by overlapping tracks across samples. **b**) Boxplots comparing the averaged DNA methylation (mCG/CG) of CGI, genic sequences (5’-UTR, CDS, intron and 3’-UTR), CREs (promoters and enhancers) and CTCF sites, and **c**) four major classes of repetitive elements between HCC and TAN. **d**) Boxplots comparing the averaged DNA methylation (mCG/CG) of hyperDMR between HCC and TAN. **e**) Genomic distribution of hyperDMR and hypoDMR in HCC. HyperDMR: promoter (*P* = 2.653e-195), UTR (*P* = 3.621e-6), exon (*P* = 0.0404); HypoDMR: intergenic (*P* = 1.1e-15). \**P* < 0.05 and \*\**P* < 0.001, one-tailed hypergeometric test corrected by the Benjamini-Hochberg approach. **f**) Boxplots comparing the expression (averaged DESeq2-normalized counts) of protein-coding genes within HCC PMDs between HCC and TAN. Number-matched, randomly selected genes were used as control. **g**) Boxplots comparing the expression (averaged DESeq2-normalized counts) of LTRs within HCC PMDs between HCC and TAN. Number-matched, randomly selected LTRs were used as control. For all boxplots, *P* values (two-tailed Wilcoxon test) are shown. The center and bounds of boxes indicate the median and quartile of all data points, respectively. The minima and maxima of whiskers indicate quartile 1−1.5×the interquartile range and quartile 3+1.5×the interquartile range, respectively.

HypoDMRs were distributed throughout the genome (**Fig. 2e**). Approximately 78% of hypoDMRs (n=100,370) overlapped with partially methylated domains (PMDs), which are large domains with lower DNA methylation as compared to the surrounding genome (**Fig. 2a** and **Extended Data Fig. 2c,d,e**)^44^. In line with previous findings, these regions were enriched with repressive histone modifications (**Extended Data Fig. 2d,e**) and were associated with transcriptional silencing (**Fig. 2f**)^45,46^. Interestingly, we found that LTR retrotransposons within PMDs, but not other classes of repeats, were depressed in HCC, despite showing lower average expression levels compared to random controls (**Fig. 2g** and **Extended Data Fig. 2f**). Overall, we uncovered widespread DNA methylation alterations at distinct genomic regions in HCC, and delineated associated changes in other epigenetic marks and transcription.

### Epigenetic alternations at promoters drives transcriptional dysregulation in HCC

Transcriptional dysregulation of the Wnt/β-catenin and TGF-β signaling pathways play key roles in mediating cell proliferation, inflammation and extracellular matrix remodeling within the HCC tumor microenvironment^6,47,48^. Indeed, we found overexpression of cell cycle regulators and suppression of immune- and extracellular matrix-related processes in our HCC samples (**Fig. 3a-c**, **Extended Data Fig. 3a** and **Supplementary Table 5**). Transcriptome-based classifications of HCC have revealed distinct clinicopathological and molecular features associated with each subclass. Of the upregulated genes that were defined as signature genes under the subclass classifications by Chiang *et al*. and Hoshida *et al*. (n=120), 78% of them (n=93) were associated with cell proliferation pathways (**Extended Data Fig. 3b,c**)^7,8^. In addition, weighted gene co-expression network analyses (WGCNA) showed that some of the dysregulated genes in our HCC samples represented the hub genes in modules that were strongly associated with HCC or TAN (**Extended Data Fig. 3d,e**). These results showed that our HCC samples exhibited transcriptomic signatures of the proliferation subclass, which comprised ∼50% of HCC^2^.

**Figure 3.**
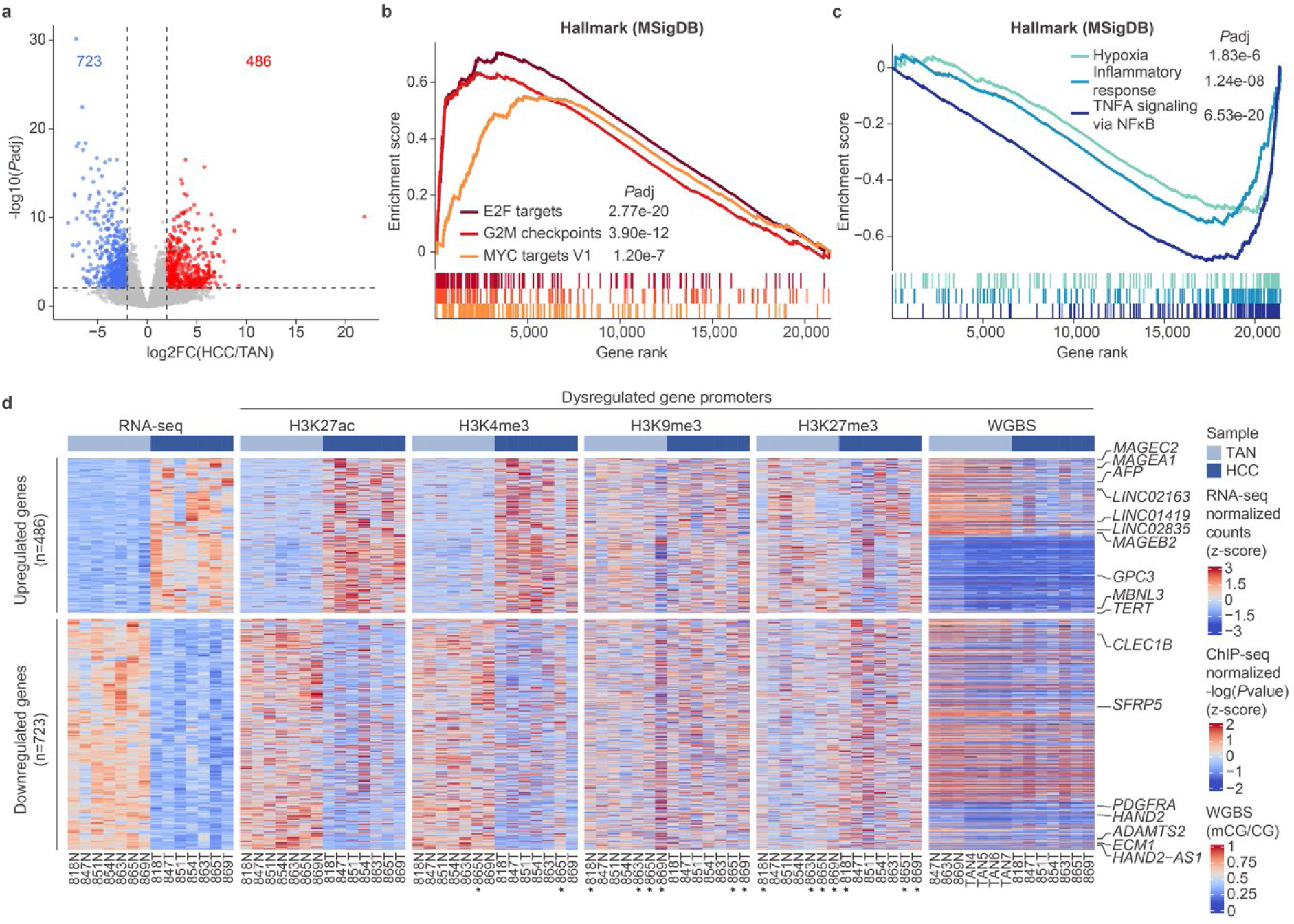
Aberrant epigenetic modifications of dysregulated genes in HCC. **a**) Volcano plot showing transcriptionally dysregulated genes in HCC. For each gene, the -log10-transformed two-tailed adjusted *P* values (*P*adj) is plotted against the log2-transformed fold change (HCC/TAN). Upregulated (*P*adj<0.01 and log2FC>1; n=486) and downregulated (*P*adj<0.01 and log2FC<-1; n=723) genes are labeled in red and blue, respectively. Dashed lines represent the indicated thresholds. **b**-**c**) GSEA revealing that dysregulated genes in HCC are positively associated with cell cycle regulation (**b**), and negatively associated with immunomodulation and extracellular matrix organization (**c**), respectively. **d**) Heatmaps showing the expression of dysregulated genes in HCC and the epigenetic signatures at their promoters in HCC and TAN samples. For each heatmap, each row represents a dysregulated gene, and each column represents a sample. Selected dysregulated genes are labeled to the right of the WGBS heatmap. Asterisks (*) indicate Avocado-imputed signals. The color scales denote the signals from RNA-seq (z-score transformed, DESeq2-normalized counts), ChIP-seq (z-score transformed - log(*P*value) of signal) or WGBS (mCG/CG).

We subsequently examined the epigenetic modifications at the promoters of dysregulated genes in our HCC samples (**Fig. 3d**). We observed concordant changes of H3K27ac and H3K4me3, but not H3K9me3 and H3K27me3, with the expression levels of dysregulated genes. Moreover, approximately half of the upregulated genes were associated with promoter DNA hypomethylation, including the HCC diagnostic marker α-fetoprotein (*AFP*), reported HCC-related lncRNAs (*LINC01419*, *LINC02163,* and *LINC02835*), and tumor-associated antigens such as the *MAGE* gene family^31–33^. Most downregulated genes were not associated with promoter DNA hypermethylation; however, 48% (n=345) overlapped with PMDs defined in HCC (**Supplementary Table 5**). For example, *CLEC1B*, a liver-enriched transmembrane glycoprotein and known prognostic marker for HCC, resided in a ∼600 kb PMD (**Extended Data Fig. 3f**)^49^. Collectively, we demonstrated an association between the transcriptional dysregulation and both local and global epigenetic changes in HCC.

### Reprogramming of typical and super enhancers perturbs gene regulatory networks in HCC

To examine the involvement of CRE reprogramming in transcriptional dysregulation in HCC, we first focused on the typical enhancers that show gain (n=3,637) or loss (n=2,924) of H3K27ac enrichment (**Fig. 4a**, **Extended Data Fig. 4a** and **Supplementary Table 6**). Notably, while the elements that lose activities were broadly shared between most HCC samples, a significant number were differentially activated amongst patients. These results suggest a degree of heterogeneity of enhancer usage in HCC. The activated typical enhancers were associated with genes related to early development (**Fig. 4a**) and were enriched for binding motifs of development-related transcription factors (TFs) such as CEBPα/β, NK2-1, and ZIC3 (**Fig. 4b**). In contrast, the inactivated typical enhancers were associated with genes relating to collagen fibril organization and chemotaxis (**Fig. 4a**) and enriched for motifs involved in biological processes including immune responses (STAT1) (**Fig. 4b**). Next, we compared the enrichment of repressive histone modifications and DNA methylation levels at the dysregulated typical enhancers in HCC and TAN (**Extended Data Fig. 4b**). While H3K27me3 enrichment did not show significant change at both the activated and inactivated typical enhancers, we observed loss of H3K9me3 enrichment at the activated typical enhancers but not in size- and number-matched, randomly selected genomic regions, suggesting that these loci may be normally repressed by H3K9me3, but not by H3K27me3. Notably, loss of DNA methylation could be detected at both the activated/inactivated typical enhancers and their controls, which could be a result of global DNA hypomethylation. Moreover, mCG is known to be positively correlated with H3K9me3 erichment^50,51^. Hence, it is possible that H3K9me3 reduction could result in mCG loss.

**Figure 4.**
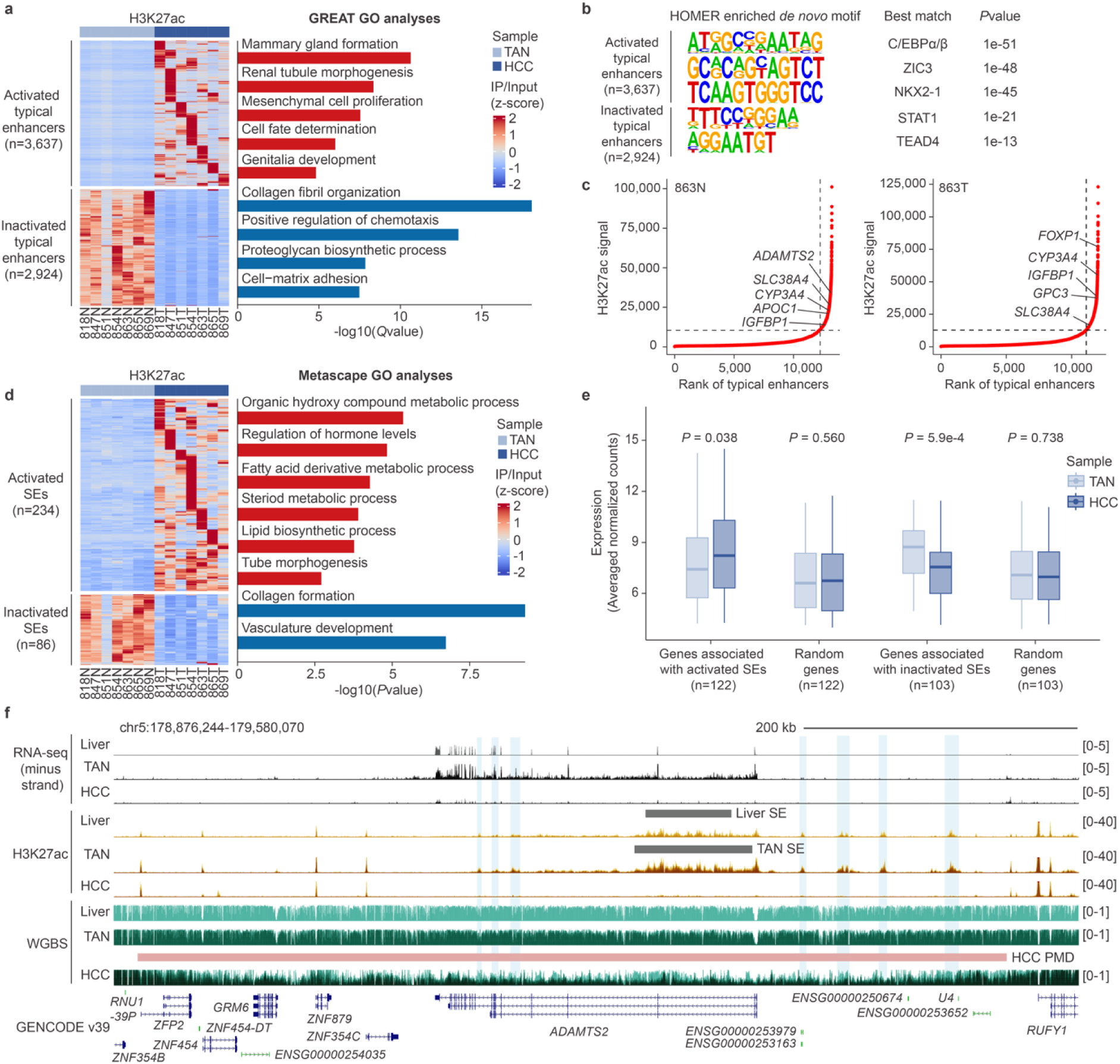
Enhancer and SE dysregulations are associated with transcriptional reprogramming in HCC. **a**) Heatmap of H3K27ac ChIP-seq signals (z-score transformed fold enrichment over input) of dysregulated typical enhancers in HCC. Each row represents a dysregulated typical enhancer, and each column represents a sample (left). Dysregulated typical enhancers in HCC were associated with GO terms related to early development and extracellular matrix remodeling (right). *Q* values (Binomial test) for the top 5 enriched GO terms are shown. **b**) Motif enrichment analyses of dysregulated typical enhancers in HCC showing enrichment of binding motifs for TFs involved in early development and immune responses. **c**) Input-normalized H3K27ac signal distribution of stitched enhancers in HCC and TAN samples from patient 863. Enhancers were ranked in increasing order based on their input-normalized H3K27ac signals by the ROSE algorithm. SEs were defined as enhancers above the inflection point of the curve. Representative genes associated with SEs are denoted. **d**) Heatmap of H3K27ac ChIP-seq signals (z-score transformed fold enrichment over input) of dysregulated putative SEs. Each row represents a dysregulated SE, and each column represents a sample (left). Dysregulated SEs were associated with metabolic and extracellular matrix-related processes (right). *P* values (one-tailed hypergeometric test) for the top 5 enriched GO terms are shown. **e**) Boxplots comparing the expression of dysregulated SE-associated genes (Averaged DESeq2-normalized counts) between HCC and TAN. Number-matched, randomly selected genes were used as control. *P* values (two-tailed Wilcoxon test) are shown. The center and bounds of boxes indicate the median and quartile of all data points, respectively. The minima and maxima of whiskers indicate quartile 1−1.5×the interquartile range and quartile 3 + 1.5 × the interquartile range, respectively. **f**) Genome browser screenshot showing an inactivated SE, and several adjacent inactivated typical enhancers located within a HCC PMD at the *ADAMT2* gene locus. The SE in liver and TAN is annotated as gray bars above their H3K27ac tracks, and the inactivated typical enhancers are highlighted by cyan shadings. The HCC PMD is annotated as pink bar above its WGBS track. Stranded total RNA-seq tracks are displayed as TPM, H3K27ac ChIP-seq tracks are displayed as fold enrichment over input, and WGBS tracks are displayed as mCG/CG. Liver RNA-seq and WGBS tracks are from Roadmap Epigenomics Project and is shown as single replicates. Other tracks are shown as composite signals by overlaying tracks across samples.

We further defined super enhancers (SEs) in our HCC and TAN samples based on the rank of H3K27ac signals of stitched typical enhancers using the ROSE algorithm (**Fig. 4c**)^52–54^. The SEs in both HCC and TAN samples were associated with metabolic genes (e.g. *CYP3A4*, *SLC38A4* and *IGFBP1*), supporting the model that SEs commonly regulate genes relevant to the cell/tissue-type functions^52–54^. We identified dysregulated SEs in HCC (n=320) (**Supplementary Table 6**). Analogous to the activated typical enhancers, the activated SEs also exhibited heterogeneity in activity across HCC samples, and were associated with genes in metabolism, early development, and extracellular matrix formation (**Fig. 4d** and **Extended Data Fig. 4c**). Importantly, we found concomitant perturbation in the expression of SE-associated genes (**Fig. 4e**). For instance, we defined an activated SE, located upstream of the *MBNL3* gene, which encodes for a splicing factor highly expressed in fetal livers and in HCC, which was also significantly upregulated in our HCC samples (**Extended Data Fig. 4d**)^55^. Conversely, downregulation of the *ADAMTS2* gene in HCC was associated with the loss of activity of a SE and several typical enhancers located within a PMD (**Fig. 4f**). Together, our results showed that the altered activities of typical enhancers and SEs were associated with transcriptional dysregulation in HCC.

### Aberrant SE reactivation concordant with *GPC3* overexpression in HCC

Our differential gene expression analyses identified the transcriptional upregulation of glypican-3 (*GPC3*) (**Fig. 3d** and **Fig. 5a**), which is another diagnostic marker often used together with AFP for HCC^36,37^. GPC3 is a heparan-sulphate proteoglycan that plays an important role in regulating cell growth and differentiation of various tissues, including the liver, during embryonic development^56^. GPC3 is highly expressed in fetal livers, silenced in adult livers, but derepressed in more than 70% of HCC tumors^37^. Due to its highly HCC-restricted expression profile (**Fig. 5b**), GPC3 has emerged as a promising immunotherapeutic target^35–37^. However, the transcriptional regulatory mechanisms for *GPC3* have not been fully explored. Interestingly, we discovered the aberrant reactivation of a SE upstream of *GPC3* in HCC (**Fig. 5c**). The aberrant SE consisted of seven constituent enhancers (E1-E7), which exhibited a similar open chromatin pattern to fetal hepatoblast and were differentially reactivated in HCC but not in other analyzed cancer types (**Fig. 5c** and **Extended Data Fig. 5a**). These constituent enhancers are bound by liver-restricted TFs (CEBPβ and HNF4A), pioneering factors (FOXA1 and FOXA2), and Wnt-signaling TFs (TCF7). Additionally, transcriptional upregulation of *GPC3* was accompanied with significant enhancer-promoter and enhancer-enhancer 3D chromatin interactions (**Fig. 5c**). We functionally validated the *cis*-regulatory potential of the seven constituent enhancers by luciferase assays and found that most candidates exhibited strong enhancer but not promoter activities (**Fig. 5d**,**e**).

**Figure 5.**
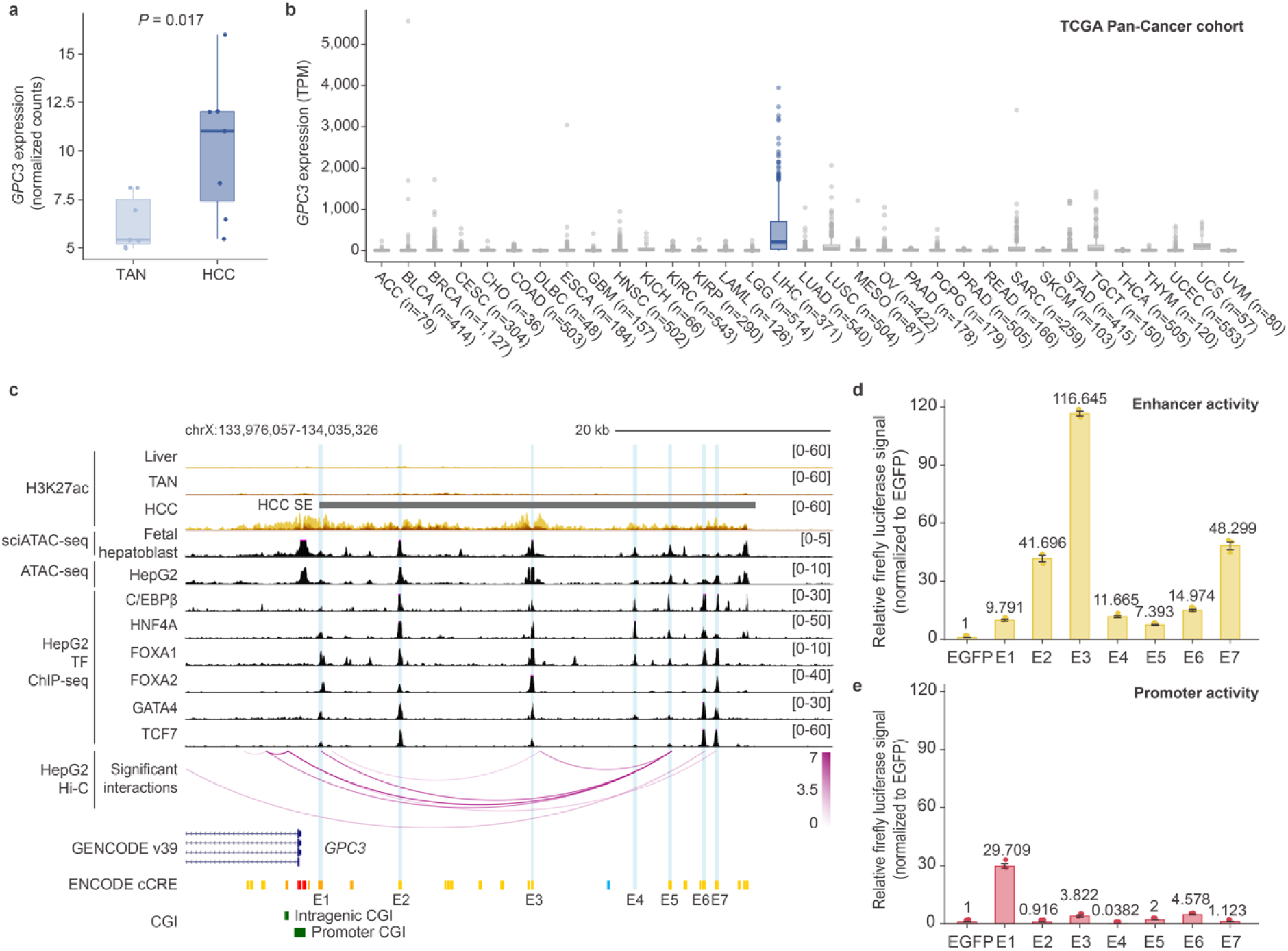
Aberrant SE reactivation drives GPC3 transcriptional upregulation in HCC. **a**) Boxplots comparing the expression of *GPC3* (DESeq2-normalized counts) between HCC and TAN. **b**) Boxplots comparing the expression of *GPC3* (TPM) across all cancer types from TCGA. HCC is highlighted in blue, while other cancer types are shown in gray. Cancer type abbreviations follow TCGA definitions. **a**-**b**) *P* values (two-tailed Wilcoxon test) are shown. The center and bounds of boxes indicate the median and quartile of all data points, respectively. The minima and maxima of whiskers indicate quartile 1−1.5×the interquartile range and quartile 3+1.5×the interquartile range, respectively. **c**) Genome browser screenshot showing the reactivated SE at the *GPC3* locus. The SE is annotated as gray bar above the HCC H3K27ac track. The constituent enhancers (E1-E7) of the *GPC3* SE are highlighted by cyan shadings. Single-cell combinatorial index-assay for transposase accessible chromatin-sequencing (sci-ATAC-seq) for fetal hepatoblast (fragment count) is from CATlas. HepG2 ATAC-seq and TF ChIP-seq (fold enrichment over input) tracks are from ENCODE. Significant interactions defined from HepG2 Hi-C data are shown as arcs and the color denotes significance (-log(*Q*value)). H3K27ac ChIP-seq tracks are displayed as fold enrichment over input. All tracks are shown as composite signals by overlaying tracks across samples. **d**-**e**) Luciferase assays assessing the enhancer (**d**) and promoter (**e**) activities of E1-E7. Data represent the mean ± standard deviation of three technical replicates from one representative experiment (n=2 independent experiments).

It should be noted that *GPC3* is also highly expressed in the placenta and is involved in placental growth and development (**Extended Data Fig. 5b**)^37,57^. We asked if placenta and HCC cells undergo similar SE reactivation. Surprisingly, we found that *GPC3* transcription in the placenta was associated with a distinct SE (**Extended Data Fig. 5c**), suggesting cell type-differential usage of SEs for regulating *GPC3*. In the TCGA-Liver Hepatocellular Carcinoma (TCGA-LIHC) cohort, we found that high expression of *GPC3* in HCC samples was not due to copy-number alterations or somatic mutations (**Extended Data Fig. 6a-c**). Furthermore, we observed no strong correlation in the expression levels of *GPC3* and the lncRNA *GPC3-AS1* (**Extended Data Fig. 6d**), which has been shown to recruit transcriptional coactivators to *GPC3* gene body to stimulate its expression^58^. Our findings suggested that aberrant reactivation of the SE contributed to transcriptional upregulation of *GPC3* in HCC.

### DNA hypomethylation of SE and CGIs modulate *GPC3* expression in HCC

Next, we investigated the epigenetic mechanisms underlying the aberrant reactivation of the *GPC3*-associated SE in HCC. We observed only subtle loss of H3K9me3 and H3K27me3 at the SE (**Extended Data Fig. 7a**). However, we detected extensive mCG loss in HCC compared to TAN at the constituent enhancers that contain two or more CpGs (**Fig. 6a**). We validated the DNA hypomethylation of these constituent enhancers in HepG2 by bisulfite amplicon sequencing (**Extended Data Fig. 7b**). Furthermore, these elements were highly methylated across normal tissues (**Fig. 6b**), suggesting a link between DNA methylation loss and the SE reactivation.

**Figure 6.**
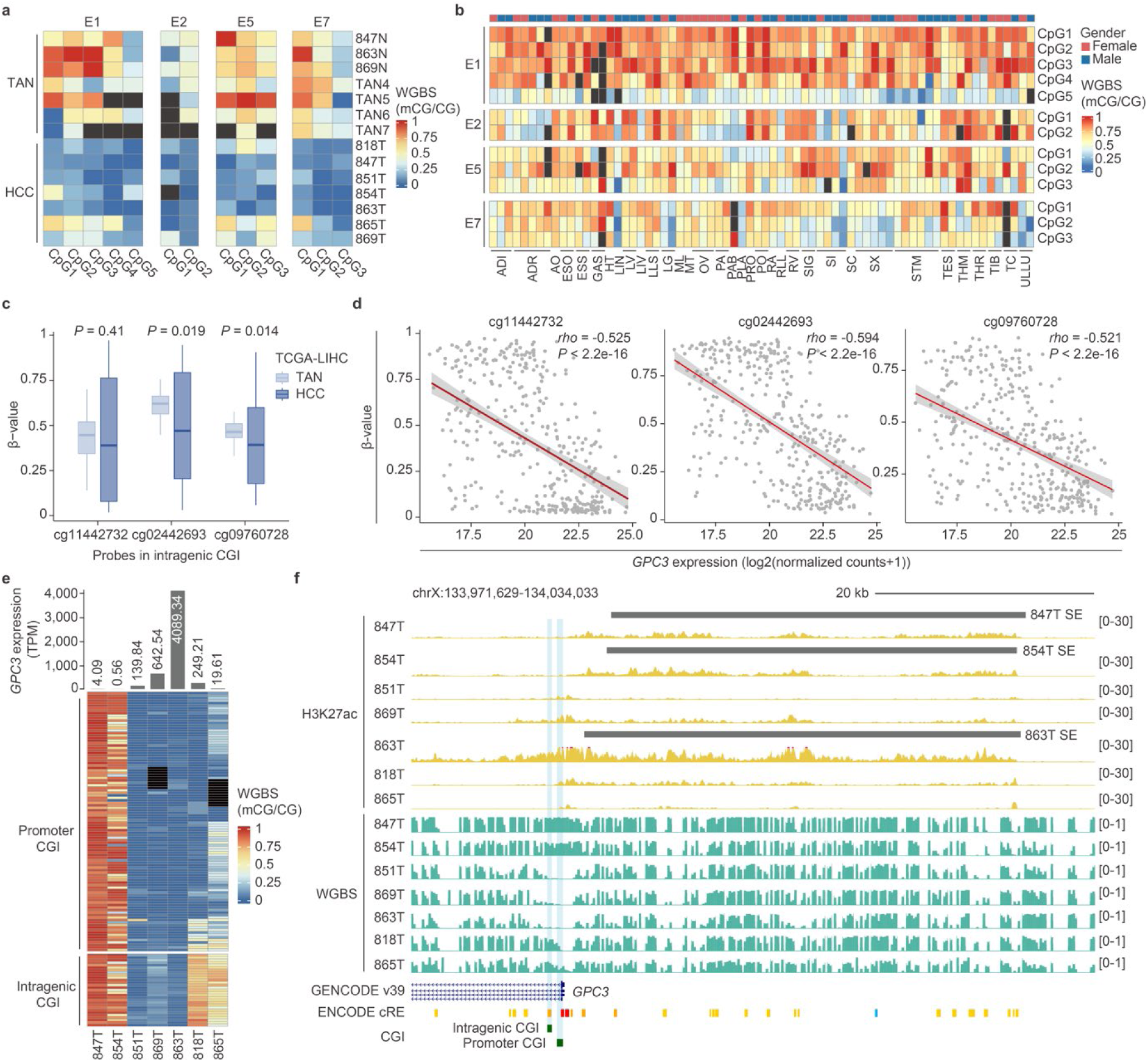
SE reactivation and DNA hypomethylation of GPC3-associated CpG islands (CGI) work in concert to regulate *GPC3* expression. **a**-**b**) Heatmap comparing the DNA methylation (mCG/CG) of constituent enhancers of *GPC3* SE that harbour at least 2 CpGs (E1, E2, E5 and E7) between HCC and TAN (**a**) and among normal tissues from ENCODE and Roadmap Epigenomics Project (**b**). **a)** Each row corresponds to a sample, and each column corresponds to a CpG within the constituent enhancer. Dark gray cells indicate CpGs with insufficient read coverage (< 5). **b**) Each row corresponds a CpG within the constituent enhancer, and each column corresponds to a sample. Dark gray cells indicate CpGs with insufficient read coverage (< 5). ADI: adipose; ADR: adrenal gland; AO: aorta; ESO: esophagus; ESS: esophagus squamous epithelium; GAS: gastroesophageal sphincter; HT: heart; LIN: large intestine; LV: heart left ventricle; LIV: liver; LLS: lower leg skin; LG: lung; ML: muscle leg; MT: muscle trunk; OV: ovary; PA: pancreas; PAB: pancreas body; PLA: placenta; PRO: prostate; PO: psoas muscle; RA: heart right atrium; SIG: sigmoid colon; SI: small intestine; SC: spinal cord; SX: spleen; STM: stomach; TES: testis; THM: thymus; THR: thyroid; TIB: tibial nerve: TC, transverse colon; ULLC: upper lobe left lung. **c**) Boxplots comparing the DNA methylation of CpGs (β-value) within the *GPC3*-associated intragenic CGI between HCC and TAN in the TCGA-LIHC cohort. *P* values (two-tailed Wilcoxon test) are shown**. d**) Scatterplots showing the correlation between the DNA methylation of CpGs (β-value) within the *GPC3*-associated intragenic CGI and *GPC3* expression (DESeq2-normalized counts) in the TCGA-LIHC cohort. Spearman’s correlation coefficient (*rho*) and *P* values are indicated at the top right of each plot. **e**) Heatmaps showing the DNA methylation (mCG/CG) of the *GPC3* promoter-associated and intragenic CGIs in each HCC sample. The corresponding *GPC3* expression (TPM) in each patient sample is shown as barplots at the top of the promoter CGI heatmap. Dark gray cells indicate CpGs with insufficient read coverage (< 5). **f**) Genome browser screenshot showing the concomitant reactivation of the *GPC3*-associated SE and DNA hypomethylation of the *GPC3* promoter-associated and intragenic CGIs. These epigenetic changes are coupled with the transcriptional upregulation of *GPC3* in HCC. SE definitions are shown as gray bars above their respective H3K27ac tracks. *GPC3* promoter-associated and intragenic CGIs are highlighted by cyan shading. H3K27ac ChIP-seq tracks are displayed as fold enrichment over input, and WGBS tracks are displayed as mCG/CG. All tracks are shown as composite signals by overlaying tracks across samples.

*GPC3* has two CGIs: one proximal to the promoter (promoter CGI) and the other within its first intron (intergenic CGI). DNA hypermethylation of the promoter CGI could silence *GPC3* transcription, but the regulatory role of the intragenic CGI has not been examined^59^. Analyzing DNA methylation microarray data from the TCGA-LIHC cohort, we found significant loss of DNA methylation at two of the CpG probes within the intragenic CGI in HCC patients (**Fig. 6c**). Importantly, the methylation levels of all CpG probes within the intragenic CGI was anticorrelated with *GPC3* expression (**Fig. 6d**). These data suggest that reduced mCG at both the promoter and the intragenic CGIs are required for *GPC3* overexpression in HCC. However, because CGIs are typically unmethylated and *GPC3* remains silenced in most healthy adult tissues, we reasoned that DNA methylation loss at the two CGIs is not sufficient to induce *GPC3* expression^60^.

Integrating our findings, we uncovered a potential relationship between the reactivation of the SE, the DNA methylation of two CGIs, and increased *GPC3* expression (**Fig. 6e**,**f**). SE reactivation without the CGIs mCG loss resulted in little to no *GPC3* expression (HCC samples 847 and 854). Similarly, DNA hypomethylation of the two CGIs but without SE reactivation resulted in relatively low *GPC3* expression (HCC samples 851 and 869). Strikingly, significant *GPC3* induction was detected in patients that had both the SE reactivation and DNA hypomethylation of the two CGIs (HCC sample 863). Importantly, DNA hypermethylation of either CGI results in lowered *GPC3* expression (HCC samples 818 and 865) and the concomitant hypermethylation of both CGIs abolishes its expression (HCC samples 847 and 854). Together, we surmised that both the aberrant reactivation of the HCC-restricted SE and DNA hypomethylation of the two CGIs work in concert to transcriptionally regulate *GPC3* in HCC.

### Aberrant retrotransposon-derived CRE activities in HCC

To characterize dysregulated retrotransposon-derived CREs in HCC, we overlapped the genomic coordinates of retrotransposons from RepeatMasker with our dysregulated CRE definitions, identifying 3,167 and 1,361 dysregulated retrotransposon-derived typical enhancers (RT-typical enhancers) and promoters (RT-promoters), respectively (**Extended Data Fig. 8a,b** and **Supplementary Table 7**). Consistent with the heterogeneity of typical enhancer/SE usage across HCC tumors (**Fig. 4a** and **Fig. 4d**), we detected differing H3K27ac enrichment levels at both dysregulated RT-typical enhancers and RT-promoters amongst patients (**Extended Data Fig. 8a,b**). The dysregulated RT-typical enhancers were also associated with GO terms relating to metabolism, early development, and extracellular matrix organization (**Fig. 4a** and **Extended Data Fig. 8c,d**). Moreover, they were enriched with developmental TF binding motifs including that of Wnt-signaling TFs (TCF3 and TCF7L1) and Hippo pathway factors (TEAD1/2/3/4) (**Extended Data Fig. 8e**). On the other hand, dysregulated RT-promoters were enriched with binding motifs for ubiquitously expressed TFs, including those for YY1, KLF, and NF-Y (**Extended Data Fig. 8e**). We next examined the class distribution of the dysregulated retrotransposon-derived CREs. Notably, we found that LTRs were over-represented in dysregulated RT-typical enhancers and RT-promoters, whereas LINEs made up a higher proportion of the dysregulated RT-typical enhancers. Analyses at the subfamily level showed that, amongst the activated RT-typical enhancers, 42 out of the 51 over-represented subfamilies were LTRs, including LTR2B, LTR8B, and LTR10A (**Supplementary Table 7**). Similarly, for the activated RT-promoters, 32 out of the 38 were LTRs subfamilies, including LTR12C and MER52A (**Supplementary Table 7**).

To further investigate the role of activated RT-promoters, we focused on those that were highly upregulated in our HCC samples (log2FC > 3.5 and *P*_adj_ < 0.01, **Supplementary Table 6**). This resulted in 24 candidate elements, of which 20 were LTRs (**Fig. 7a**). Interestingly, 11 of these LTRs overlapped with lncRNA promoters (**Fig. 7a**). The activation of these elements was associated with loss of repressive epigenetic signatures (**Fig. 7a**), suggesting that epigenetic reprogramming could contribute to their transcriptional derepression in HCC.

**Figure 7.**
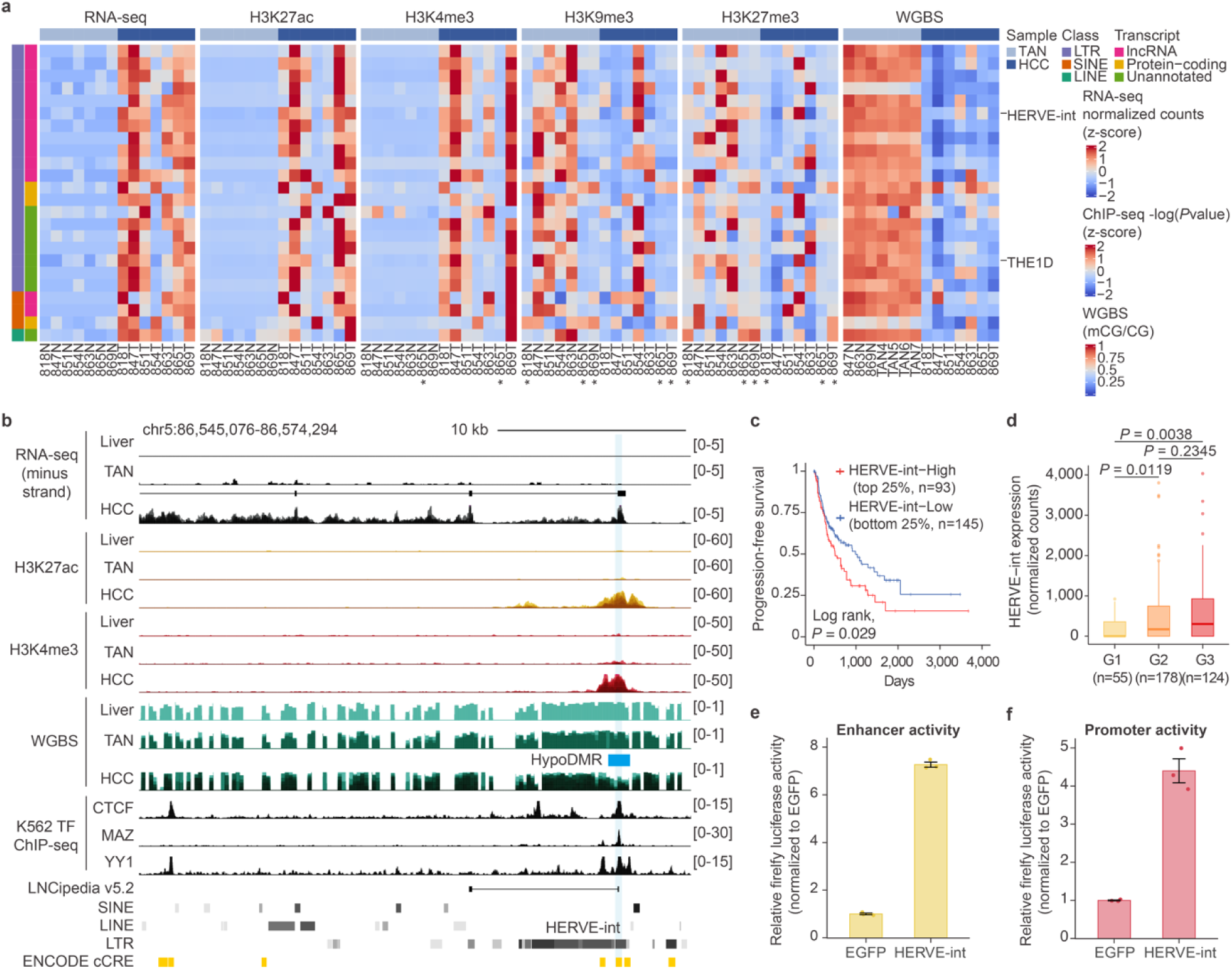
Epigenetic dysregulation of retrotransposon-driven transcripts in HCC. **a**) Heatmap showing the expression and epigenetic signatures of significantly dysregulated activated RT-promoters (n=24) in HCC. Each row represents a dysregulated element, and each column represents a sample. The color scales denote the signals from RNA-seq (z-score transformed, DESeq2-normalized counts), ChIP-seq (z-score transformed -log(*P*value) of signal), or WGBS (mCG/CG). **b**) Genome browser screenshot showing the expression and epigenetic signatures of HERVE-int. StringTie-assembled transcripts are shown above the HCC RNA-seq track. HERVE-int is highlighted by cyan shading. HypoDMR in HCC is annotated as blue bar above the WGBSs track. K562 TF ChIP-seq tracks (fold enrichment over input) are from ENCODE. Stranded total RNA-seq tracks are displayed as TPM, ChIP-seq tracks are displayed as fold enrichment over input, and WGBS tracks are displayed as mCG/CG. Liver RNA-seq and WGBS tracks are from Roadmap Epigenomics Project and shown as single replicates. Other tracks are shown as composite signals by overlaying tracks across samples. **c**) Kaplan-Meier survival curve for HERVE-int expression in the TCGA-LIHC cohort. Based on HERVE-int expression (DESeq2-normalized counts), patients were stratified into the low (bottom 25%) and high (top 25%). *P* value (log-rank test) is shown. **d**) Boxplot comparing the expression of HERVE-int (DESeq2-normalized counts) across tumor grades in the TCGA-LIHC cohort. *P* values (two-tailed Wilcoxon test) are shown. **e**-**f**) Luciferase assays assessing the enhancer (**e**) and promoter (**f**) activities of HERVE-int. Data represent the mean ± standard deviation of three technical replicates from one representative experiment (n=2 independent experiments).

### Global DNA hypomethylation leads to cryptic activation of LTR retrotransposons with clinical relevance in HCC

Next, we analyzed the expression patterns of the retrotransposon-driven transcripts amongst HCC patients from the TCGA-LIHC cohort. Patients were stratified based on the expression levels of each of the 24 elements. Strikingly, we discovered that the expression of two elements, belonging to the HERVE (HERVE-int) and THE1D subfamilies, could significantly differentiate patients’ survival (**Fig. 7b**,**c** and **Extended Data Fig. 9a,b**). Patients with high HERVE-int expression (HERVE-int-High, top25%, n=93) were associated with poorer progression-free survival (**Fig. 7c**). HERVE-int expression also increased across tumor grades, supporting its prognostic potential (**Fig. 7d**). Similarly, patients with intermediate to high THE1D expression (THE1D-Intermediate, 30%<x<70%, n=149; THE1D-High, top30%, n=112) also exhibited significantly poorer progression-free survival (**Extended Data Fig. 9c)**. It should be noted that different stratifying criteria were applied for HERVE-int and THE1D due to their distinct distribution of expression patterns amongst HCC patients (**Supplementary Table 8**).

The HERVE-int element overlapped with a lncRNA promoter and was transcribed in the negative strand orientation (Fig. 7b), whereas the THE1D element served as the alternative promoter for the pseudogene *ENSG00000286540* encoded on the positive strand (**Extended Data Fig. 9a,b**). Transcriptional activation of both elements in HCC was associated with loss of DNA methylation (**Fig. 7b**, **Extended Data Fig. 9a,b,d** and **Extended Data Fig. 10a**). To test the mechanistic relationship between the retrotransposon activities and DNA methylation loss, we treated HCC cell line models with the DNA demethylating agent 5-aza-2’-deoxycytidine (DAC). Given that THE1D was expressed in our HCC cell lines, we focused on analyzing HERVE-int expression change in HepG2 cells, which showed no HERVE-int activity (**Extended Data Fig. 10b**). DAC-treatment resulted in 50-70% decrease in global DNA methylation (**Extended Data Fig. 10c**) concomitant with ∼35-fold induction in HERVE-int expression (**Extended Data Fig. 10d**). These results suggest that mCG loss was sufficient for HERVE-int reactivation. Given the potential for LTRs to serve as CREs, we performed luciferase assays to demonstrate that the HERVE-int element could function both as an enhancer and a promoter (**Fig. 7e**,**f**). Furthermore, analyzing publicly available TF ChIP-seq data, we discovered that HERVE-int could be targeted by CTCF and YY1, both of which are involved in mediating enhancer-promoter interactions (**Fig. 7b**)^61–63^. Thus, loss of DNA methylation at HERVE-int could facilitate CTCF and YY1 binding, leading to aberrant *cis*-regulatory interactions.

To elucidate the regulatory role of HERVE-int and THE1D in HCC, we performed CRISPR interference (CRISPRi) to transcriptionally repress the two elements. We successfully depleted HERVE-int and THE1D expression by approximately 80-90% (**Fig. 8a** and **Extended Data Fig. 9e**). Next, we generated RNA-seq datasets from the cells depleted of either HERVE-int or THE1D. We found that silencing of either element did not have wide-spread impact transcription (**Fig. 8b**, **Extended Data Fig. 9f** and **Supplementary Table 9**). However, specific genes with reported roles in cancers were significantly dysregulated. For instance, in the HERVE-int-depleted cells, *ANXA1*, which has anti-inflammatory activities in various cancers, and *DUSP9*, which is highly expressed in less differentiated HCC tumors, were upregulated and downregulated, respectively (**Fig. 8b**)^64,65^. In the THE1D-depleted cells, *RBP1*, a gene downregulated in > 80% of HCC, was upregulated (**Extended Data Fig. 9f**)^66^.

**Figure 8.**
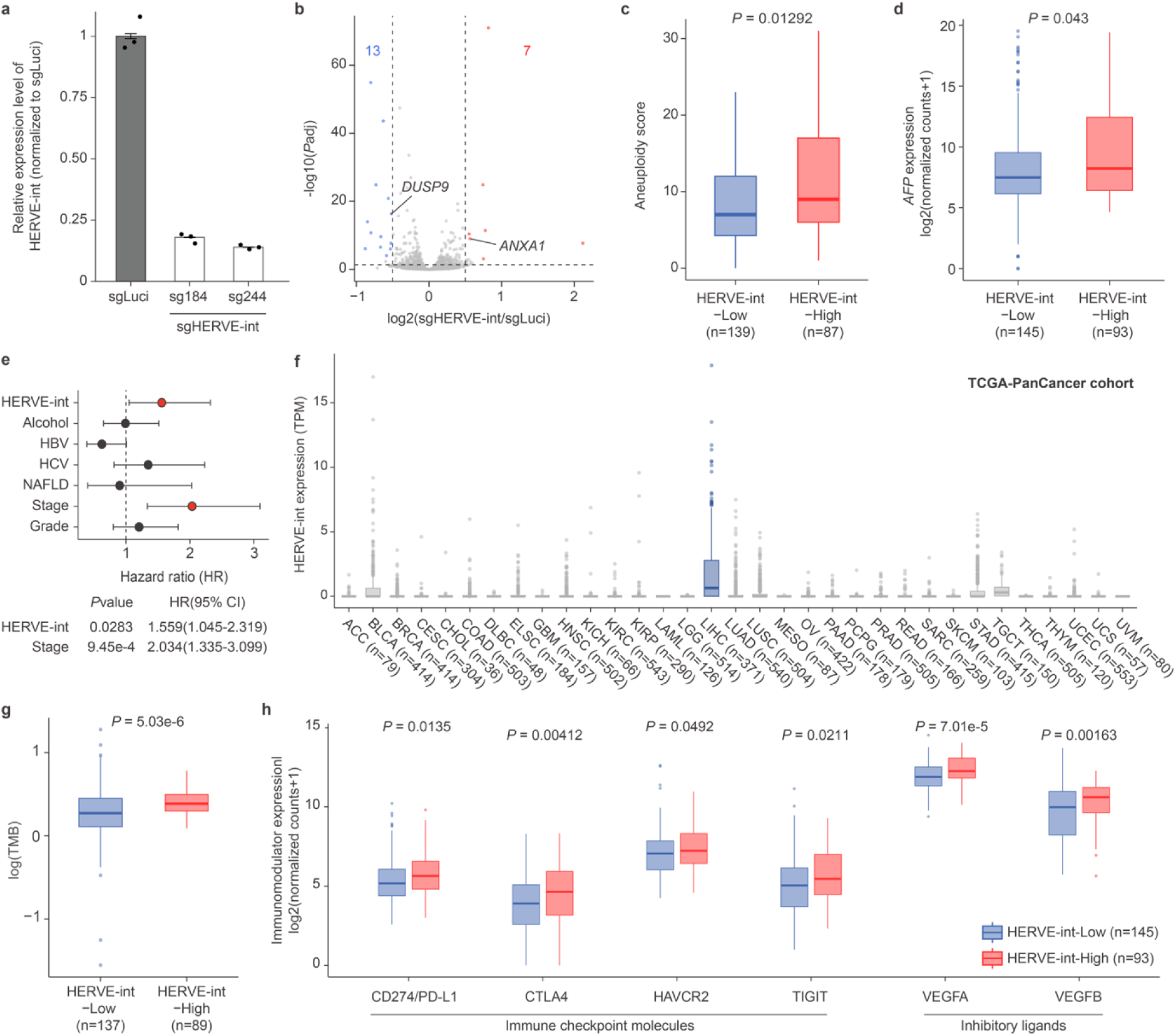
HERVE-int dysregulation in HCC is associated with specific molecular signatures in HCC and might serve as potential indicator of better response to immunotherapy. **a**) Barplots showing the relative expression of HERVE-int measured by RT-qPCR in PLC cells transduced with HERVE-int-targeting sgRNAs (sgHERVE-int) or luciferase-targeting sgRNA control (sgLuci). Each dot represents a technical replicate and error bars denote standard deviations. *GAPDH* is used as internal control and HERVE-int expression is normalized to PLC cells transduced with sgLuci. **b**) Volcano plot showing dysregulated genes in PLC cells depleted of HERVE-int. For each gene, the -log10-transformed, two-tailed adjusted *P* values (*P*adj) is plotted against the log2-transformed fold change (HCC/TAN). Upregulated (*P*adj < 0.05 and log2(Fold change (sgHERVE-int/sgLuci) > 0.5) and downregulated (*P*adj < 0.05 and log2(Fold change (sgHERVE-int//sgLuci) < -0.5) are labeled in red and blue, respectively. Dashed lines represent the indicated thresholds. Selected dysregulated genes are labeled. **c**-**d**) Boxplots comparing the aneuploidy score (**c**) and *AFP* expression (log2-transformed, DESeq2-normalized counts) (**d**) between HERVE-int-Low and HERVE-int-High patients. **c**) Only patients with matched aneuploidy score were analyzed. **e**) Multivariate cox regression analyses for HERVE-int expression and other prognostic factors for HCC patients in the TCGA-LIHC cohort. Patients with low HERVE-int expression, no history of alcohol consumption, no HBV/HCV infection, no non-alcoholic fatty liver disease (NAFLD), early tumor states (I/II) or low tumor grades (G1/2) were used as the reference level for the respective prognostic factor in the model. *P* values (log-rank test) and 95% confidence intervals for hazard ratios of the two independent prognostic indicators of the model are shown. **f**) Boxplots comparing the expression of HERVE-int (TPM) across all cancer types from TCGA. HCC is highlighted in blue, while other cancer types are shown in gray. Cancer type abbreviations follow TCGA definitions. **g**) Boxplots comparing the log10-transformed TMB between HERVE-int-Low and HERVE-int-High patients. Only patients with matched mutation data were analyzed. **h**) Boxplots comparing the expressions (log2-transformed, DESeq2-normalized counts) of selected immune checkpoint molecules and inhibitory ligands between HERVE-int-Low and HERVE-int-High patients. For all boxplots, *P* values (two-tailed Wilcoxon test) are shown. The center and bounds of boxes indicate the median and quartile of all data points, respectively. The minima and maxima of whiskers indicate quartile 1−1.5×the interquartile range and quartile 3+1.5×the interquartile range, respectively.

We then assessed whether HCC patients showing increased expression of HERVE-int or THE1D harbored molecular characteristics that were linked to their worse disease outcomes. Previous multi-omic characterization had established the iCluster1-3 molecular classifications for HCC^6^. Using these definitions, we found that both HERVE-int-High and THE1D-High patients among the TCGA-LIHC cohort were strongly associated with the iCluster3 group (**Extended Data Fig. 9g**). This subclass is characterized by increased chromosomal instability and increased expression of the *AFP* marker gene^6^. HERVE-int-High, but not THE1D-Intermediate or -High patients, showed higher incidences of these molecular phenotypes (**Fig. 8c**,**d**). In line with the more aggressive tumors of the iCluster3 subclass, we discovered that high HERVE-int expression could serve as an independent prognostic factor for HCC (**Fig. 8e**). Moreover, HERVE-int exhibited higher expression in HCC compared to other cancer types (**Fig. 8f**). Also, consistent with the increased chromosomal instability, HERVE-int-High patients exhibited higher tumor mutational burden (TMB) (**Fig. 8g**)

In addition to the accumulation of molecular aberrations, the immune microenvironment also plays a key role in tumor development and progression. Intriguingly, we observed upregulation of key immunomodulators, including those encoding for the immune checkpoint molecules PD-L1 and CTLA4, and VEGF ligands, which are important angiogenesis mediators that could blunt anti-tumor immune responses (**Fig. 8h**)^67–69^. While these findings hint at the altered immune landscapes of HERVE-int-High patients, further studies are needed to delineate the precise consequences of these changes. Collectively, these results point to the clinical relevance of epigenetic dysregulation of retrotransposons and present HERVE-int as potential prognostic indicators for HCC.

## DISCUSSION

In this study, we performed integrated epigenomic and transcriptomic characterization of HCC. We leveraged the enriched datasets to elucidate the transcriptional regulatory landscape. Our work builds upon previous epigenomic studies of HCC by demonstrating the impact of epigenetic changes on CRE and retrotransposon activities at both local and global levels. At a locus-specific level, we showed the transcriptional changes that arose from focal DNA hypomethylation of promoters. At the global level, we found that mCG loss at PMDs, coupled with enrichment of repressive histone modifications, contribute to extensive transcriptional inhibition.

Transcriptional dysregulation in cancers can stimulate pathways that lead to dedifferentiation and loss of cellular identity. Such processes are driven, in part, through reactivation of genes that confer cancer stemness. Indeed, dysregulated typical enhancers and SEs in HCC were associated with developmental processes. In other cancer types, BET-bromodomain inhibitor JQ1 has been used to selectively disrupt SEs that regulate oncogenes^53^. Our data supports further exploration of SE targeting as a therapeutic potential in HCC. Furthermore, we discovered the aberrant reactivation of a fetal liver SE associated with the oncofetal protein GPC3^37^. GPC3 is a diagnostic marker and immunotherapeutic target for HCC^35–38^. While anti-GPC3 antibodies and anti-GPC3 chimeric antigen receptor (CAR) T-cell therapy have been developed to target GPC3 in HCC, they have achieved mixed clinical success due to variable response rates^35,38^. Earlier studies showed that DNA hypomethylation of GPC3 promoter CGI was insufficient to account for the differential expression of *GPC3* in HCC^59^. Here, our findings demonstrate an additional epigenetic mode of *GPC3* transcriptional regulation. We provided evidence that aberrant reactivation of an upstream SE and loss of DNA methylation at two *GPC3*-associated CGIs are the prerequisites for high *GPC3* expression in HCC. Given that GPC3-negative HCC patients could be characterized by having DNA hypermethylation of the GPC3 promoter and/or intragenic CGIs despite the aberrant SE reactivation, our results suggest potential value of combination therapies that include DAC and anti-GPC3 immunotherapy to sensitize GPC3-negative HCC tumors. Moreover, DAC-treatment will likely also increase immunogenicity of tumors via the viral mimicry phenomenon^70,71^. Nevertheless, further investigations are warranted to investigate this potential in greater depth.

Global DNA hypomethylation could result in activation of retrotransposon-derived CREs, disrupt gene regulatory networks, and impact cancer cell fitness^22,24–26,29^. Consistent with this, we observed aberrant retrotransposon-derived CRE activities in our HCC samples. Moreover, we showed that global DNA hypomethylation could lead to transcriptional derepression of specific elements with prognostic potentials. Specifically, we found that HCC patients with high expression of two LTRs, a HERVE-int element and a THE1D element, were associated with poorer disease outcomes. These patients also exhibited specific molecular signatures associated with advanced tumor stages. Several lines of evidence have demonstrated a connection between retrotransposon expression and anti-tumor immune responses^24,70–75^. For instance, increased ERV activities have been reported in clear cell renal cell carcinoma, which has relatively low TMB but are highly immunogenic^76,77^. Moreover, the expression of these elements was positively associated with clinical response to anti-PD-1 inhibitor^76^. Intriguingly, we found that HERVE-int-High HCC patients had higher expression of key immune checkpoint molecules and VEGFA/B, suggesting that this subgroup of patients could be more responsive to immune checkpoint blockade therapy and/or combined therapy with anti-VEGF inhibitor. In recent years, the combination of Atezolizumab (anti-PD-L1) and bevacizumab (anti-VEGF) treatments has demonstrated more potent antitumor activities and have become the new standard of care for patients with unresectable HCC^78,79^. While high TMB and high expression of immune checkpoint molecules, including PD-L1, have emerged as potential predictive biomarkers for immunotherapy responsiveness across cancer types, they have limited clinical utility when used in isolation^80–83^. Importantly, there is currently no validated predictive biomarkers for HCC^3^. Integrating additional molecular signatures, such as the expression of specific retrotransposons into the predictive marker panels could enhance the identification of HCC patients most likely to benefit from immunotherapy. Taken together, our results not only highlight retrotransposons as potential prognostic indicators, but also shed light on their possible contribution to predictive signatures for immunotherapy response in HCC.

In summary, our studies provide an integrative analysis of the transcriptional regulatory landscape of HCC and the involvement of epigenetic dysregulation of CREs and retrotransposons in hepatocarcinogenesis. Future studies examining etiology- and subtype-specific epigenetic changes could lead to improved precision in clinical management of HCC.

## Acknowledgements

We thank Prof. Nathalie Wong for sharing the PLC cell line, Dr. Gary Hon for sharing the Lenti-dCas9-KRAB-blast plasmid, and all staff in the HKUST Biosciences Central Research Facility for their assistance in carrying out the study. We also acknowledge the Centre for PanorOmic Sciences (CPOS) at the University of Hong Kong for generating a subset of the HCC WGBS data.

## Author contributions

C.C.Y.C., K.Y.Y., A.S.C., and D.L. conceived the study. P.B.S.L., and W.Y. collected and processed the clinical samples. M.F.C., A.Y.L., C.C.Y.C., G.L., and J.Y.J.A. performed all the experiments. C.C.Y.C., I.R.M., M.F.C., Q.W., H.W., and S.C.H.C. conducted all data analyses. C.C.Y.C., K.Y.Y., A.S.C., and D.L. prepared the manuscript with input from all authors.

## Supplementary Materials

Extended Data Figures 1-10

Supplementary Table 1

Legends for Supplementary Tables 1-10

## MATERIALS AND METHODS

### Patient consent and specimen processing

Seven HBV-associated HCC patients who underwent hepatectomy at the Prince of Wales Hospital (Hong Kong) were enrolled in this study. Informed consent was obtained from all patients, and the study protocol was approved by the Joint Chinese University of Hong Kong-New Territories East Cluster Clinical Research Ethics Committee. Patient clinical metadata is summarized in **Supplementary Table 1**. Patient specimens were processed immediately after surgery and snap-frozen in liquid nitrogen for storage. Frozen liver tissues were pulverized with mortar and pestle in liquid for genomic DNA and RNA extraction.

### Cell culture

HepG2 was purchased from ATCC (ATCC HB-8065). PLC was a gift from Prof. Nathalie Wong (Department of Surgery, The Chinese University of Hong Kong). HCC cells were maintained in Dulbecco’s modified Eagle’s Medium (DMEM) supplemented with 10% fetal bovine serum (Gibco), 1% L-glutamine (Gibco), 1% non-essential amino acids (Gibco), 1% sodium pyruvate (Gibco) and 1% penicillin-streptomycin (Gibco) at 37*°C* in 5% CO_2_. Medium was changed every 48 hours and cells were passed once they reach 80% confluency.

### Whole genome bisulfite sequencing

To mitigate amplification bias, four WGBS libraries were prepared per sample. WGBS libraries for the seven HCC tissues were prepared by the Centre for PanorOmic Sciences, The University of Hong Kong. Briefly, DNA was sonicated to 200-400 bp fragment. 100 ng of the fragmented DNA was spiked with 1% lambda DNA (Promega) and subjected to bisulfite conversion using EZ DNA Methylation Lighting Kit (Zymo Research Corp., Orange, CA, USA). Libraries were prepared using the EpiGenome™ Methyl-Seq Kit following manufacturer’s instructions. Each library was sequenced on a different lane on the Illumina HiSeq1500 platform. WGBS libraries for the three TAN tissues were prepared at the Center for Epigenomics Research, The Hong Kong University of Science and Technology. Briefly, genomic DNA was extracted using DNeasy® Blood & Tissue Kit (Qiagen) following manufacturer’s protocol. 1 µg of genomic DNA spiked with 0.5% lambda DNA (Promega) was sonicated to 100-1500 bp using Covaris® S200 focused-ultrasonicator (Covaris). Adaptor-ligated DNA fragments (200-400 bp) were bisulfite converted using EpiJET Bisulfite Conversion Kit (Thermo Fisher Scientific) and amplified using KAPA HiFi HotStart Uracil+ReadyMix (KAPA) for 7 cycles. Libraries were sequenced on Illumina HiSeq 4000. WGBS quality metrices are listed in **Supplementary Table 2**.

### ChIP-seq

MicroChIP for patient samples were performed as previously described^84^. Briefly, pulverized liver tissues were crosslinked with 1% formaldehyde (Sigma-Aldrich) in 1×PBS for 8 minutes at room temperature, and then quenched with glycine at a final concentration of 125 mM for 5 minutes at room temperature. After washing with ice-cold 1 ×PBS twice, pulverized and crosslinked liver tissues were sonicated using Covaris® S200 focused-ultrasonicator (Covaris), and fragmented chromatin was extracted by RIPA dilution buffer. Protein A Dynabeads (Thermo Fisher Scientific) were bound with antibodies of interests (H3K27ac, Active Motif 39133; H3K4me3, Active Motif 39915; H3K9me3, Abcam ab176916 or H3K27me3, Active Motif 39155) and incubated at 4*°*C for 36 hours. Captured chromatin was washed four times with RIPA buffer and eluted at 37°C for 1 hour in ChIP elution buffer. Eluted chromatin was crosslinked by incubating with Proteinase K (NEB) at 68°C for 4 hours, and DNA was purified with QIAquick*®* PCR Purification Kit (Qiagen). Libraries were prepared using KAPA HyperPrep DNA Kit (Roche) following manufacturer’s instruction and sequenced on Illumina NextSeq 500.

Native ChIP for HCC cells were performed as previously described with minor modifications^85^. Briefly, 1×10^7^ cells were resuspended in ice-cold douncing buffer and lysed by passing through 25GX 5/8*"* needle (Terumo) for 50 times. Chromatin was digested with 550U/mL MNase (Thermo Fisher Scientific) for 15 minutes at 37°C without shaking. EDTA was added to quench the digestion. Nuclear membrane was disrupted by incubating 1 hour in hypotonic lysis buffer. ChIP was performed by incubating digested chromatin with antibodies that were prebound to Dynabeads*™* M-280 Sheep Anti-Mouse IgG (Thermo Fisher Scientific) or Dynabeads*™* M-280 Sheep Anti-Rabbit IgG (Thermo Fisher Scientific) in IP buffer overnight at 4°C. Captured chromatin was washed twice with wash buffer, once with final wash buffer, and eluted at 68°C for 2 hours in elution buffer together with RNase A (Sigma-Aldrich). Eluted DNA was purified by QIAquick*®* PCR purification Kit (Qiagen) following manufacturer’s instruction and sequenced on Illumina NextSeq 500. ChIP-seq quality metrices are listed in **Supplementary Table 2**.

### RNA-seq and RT-qPCR

For stranded total RNA-seq, total RNA was extracted using RNeasy® Mini Kit (Qiagen). 200 ng (patient samples) or 1 µg (HCC cell lines) of total RNA was subjected to rRNA depletion using Ribo-off rRNA depletion Kit (H/M/R) (Vazyme), followed by library preparation using QIAseq™ Stranded Library Kit (Qiagen) as per instruction. Libraries were sequenced on Illumina NextSeq 500.

For RT-qPCR, total RNA was extracted using RNeasy® Mini Kit (Qiagen). 1 µg of total RNA was treated with DNaseI (NEB) and purified with Agencourt® RNAClean™ XP (Beckman Coulter). First-strand synthesis was performed using SuperScript® III Reverse Transcriptase (Invitrogen) following manufacturer’s protocol. cDNA templates were subjected to 40 cycles of PCR using LightCycler 480 Instrument II (Roche). RT-qPCR primers used are listed in **Supplementary Table 3**.

### Luciferase assays

Luciferase assays for the constituent enhancers of the *GPC3* SE and HERVE-int were performed in HepG2 and PLC cells, respectively. Briefly, the candidates were cloned downstream of the luciferase reporter gene in pGL3-promoter vector (Promega) to test for enhancer activity, or upstream of the luciferase reporter gene in pGL3-enhancer vector (Promega) to test for promoter activity, respectively. A random EGFP sequence that has no *cis*-regulatory activity was also cloned accordingly and used as negative controls. 0.8×10^5^ cells were seeded per well in 24-well plate 18 hours prior experiment. The cells were co-transfected with pGL3 vector containing the candidate or random EGFP sequence and pGL4.75[hRluc/CMV] vector (Promega) in 1000:1 pmol ratio using Lipofectamine™ 3000 Transfection Reagent (Thermo Fisher Scientific). Luciferase signals were measured 48 hours after transfection using Dual-Luciferase® Reporter (DLR™) Assay System as per manufacturer’s protocol on a FlexStation3® Multi-Mode Microplate Reader. Enhancer or promoter activity of each candidate was calculated using the firefly-to-renilla signal ratio and normalized to that of the random EGFP sequencing. Primers used for cloning are listed in **Supplementary Table 3**.

### Bisulfite amplicon sequencing for constituent enhancers of the *GPC3* SE

HepG2 genomic DNA was extracted using Allprep® DNA/RNA Mini Kit (Qiagen). 500 ng of genomic DNA was bisulfite converted using EpiJET Bisulfite Conversion Kit (Thermo Fisher Scientific) following manufacturer’s protocol. PCR was performed using TaKaRa EpiTaq™ HS (Takara Bio Inc.) with sequence specific primers. Fragments of interest were TA cloned (Thermo Fisher Scientific) and Sanger sequenced. Lollipop diagrams were generated using BiQ Analyzer^86^. Bisulfite PCR primer used are listed in **Supplementary Table 3**.

### DAC treatment and HERVE-int RT-qPCR

1×10^6^ HepG2 cells were seeded per well in 6-well plate 18 hours prior treatment. HepG2 cells were treated with 10 µM DAC (Sigma-Aldrich) or DMSO (Sigma-Aldrich) for 48 hours, with replenishment at 24 hours post treatment. DAC was then washed out, and cells were allowed 48 hours for recovery before harvesting for HERVE-int RT-qPCR.

### CRISPRi knockdown of HERVE-int and THE1D

CRISPRi for HERVE-int and THE1D was performed on PLC-dCas9 and HepG2-dCas9 cells, respectively. The dCas9 stable cells were generated as previously described with minor modifications^87^. Briefly, HEK293 cells were seeded at 1.2×10^6^ cells per well of 6-well plate and transfected with pMD2.G (Addgene plasmid # 12259), pCMVR8.74 (Addgene plasmid # 22036) and Lenti-dCas9-KRAB-blast (Addgene plasmid # 89567) using Lipofectamine™ 3000 Transfection Reagent (Thermo Fisher Scientific). Cells were incubated at 37*°*C in 5% CO_2_ overnight and media was changed after 18 hours. Viral supernatant was harvested at day 2 and 3 post-transfection and were used to infect cells with 8 µg/mL filter-sterilized polybrene. Successfully transduced cells were selected with 10 µg/mL blasticidin (Invivogen).

sgRNAs targeting HERVE-int, THE1D or luciferase gene were designed using CRISPOR v5.0.1 and cloned into pLKO5.sgRNA.EFS.GFP (Addgene plasmid # 57822)^88^. CRISPRi experiments were performed following procedures as described above by including the pLKO5.sgRNA.EGFP vector carrying sgRNA targeting HERVE-int, THE1D or luciferase gene. GFP-positive cells were FACS sorted using BD FACSAria™ III (BD Biosciences). RNA-seq libraries were generated as described above. sgRNA oligos are listed in **Supplementary Table 3**.

### Avocado imputation

Avocado v0.4.0 was used for model training and signal imputation^40^. keras v2.13.1 library was used in conjunction with CUDA v11.8 and cuDNN v8.8.0.121 to interface with TensorFlow v2.13.0 library for all model training. -log10(*P*values) signals from each chromosome were binned at a 25 bp resolution and arcsinh transformed. Models were trained on a per-chromosome basis with parameters: –batch_size=40000 –n_celltype_factors=32 – n_assay_factors=256 –n_25bp_factors=25 –n_250bp_factors=40 –n_5kbp_factors=45 – n_layers=2 –n_nodes=2048 and –n_epoch2=100. The ADAM optimizer and mean-square error (MSE) loss were employed for fitting the in-house data to the Avocado ENCODE-Core hg38 pre-trained model (https://zenodo.org/records/4774521) using the fit_celltypes and fit functions. The epoch size was adjusted for each chromosome by dividing the number of genomic positions (chromosome length divided by a bin size of 25) by a batch size of 40000 and rounding to the nearest integer. In the evaluation phase, 10 out of 43 samples were left-out from the training set to assess the per-sample global MSE of the imputed signal (mean of (imputed bin values−ground truth bin values)^2). This evaluation was compared against two baseline measures: (1) the per-sample baseline MSE (mean across all bins (ground truth value of one bin of that sample - mean of all ground truth bin values of that sample)^2), and 2) baseline MSE averaged across all other samples (mean across all bins (ground truth value of one bin of that sample - mean of all ground truth bin values across all other samples)^2).

### WGBS data analyses

#### Mapping and visualization

Paired-end sequencing reads were aligned to GRCh38/hg38 using Bismark v0.23.0 with default parameters^89^. PCR duplicates were removed by Picard tools v2.23.0 (http://broadinstitute.github.io/picard/). Only CpGs with at least 5 reads were retained for all downstream analyses. Signal tracks with mCG/CG per 200 bp bin were generated for visualization in genome browser. Circos plot comparing the DNA methylation landscape of HCC and TAN was generated using pyCirclize v1.3.0 (https://github.com/moshi4/pyCirclize) in python 3.12.

#### Identification of DMRs

DMRs were called with methylkit v1.20.0 per 500 bp non-overlapping window with at least 5 CpGs covered^90^. Significance of %mCG difference in each window was determined by Fisher’s exact test with *Q*value adjusted by SLIM method^91^. HyperDMRs in HCC were defined as regions with at least two CpGs showing ≥ 30% gain in mCG. Similarly, hypoDMRs in HCC were defined as regions with at least two CpGs showing ≥ 30% loss in mCG. HyperDMRs and hypoDMRs in HCC are listed in **Supplementary Table 4**.

#### Identification of PMDs

PMDs were defined as previously described^92,93^. Briefly, PMDs were called with methpipe v5.0.1 using default parameters^94^. Only PMDs with sizes no less than 50 kb, and do not overlap with gaps in GRCh38/hg38 or the HOX clusters were retained for analyses. For protein-coding gene or repeat expression quantification within PMDs, only those with at least 80% of length overlapping with PMDs were considered. PMDs in HCC are listed in **Supplementary Table 4**.

### RNA-seq data analyses

#### Mapping and visualization

Paired-end sequencing reads were aligned to GRCh38/hg38 and GENCODE v39 with STAR v2.7.6 using ENCODE parameters except where –outFilterMultimapNmax was set to 1 to keep only uniquely mapped reads^95^. TPM of genes were quantified with RSEM v1.3.3^96^. TPM signal tracks were generated for visualization in genome browser, using the bamCoverage program in deepTools v3.5.1 with parameters –normalizeUsing BPM –exactScaling and – filterRNAstrand forward/reverse for minus/plus strand, respectively^97^.

#### Identification of dysregulated genes and retrotransposons

Only transcriptionally active genes (TPM>1 in ≥2 samples) and retrotransposons (RPKM>1 in ≥ 3 samples) were used as inputs for differential analyses. Dysregulated genes and retrotransposons were defined using DESeq2 v1.36.0^98^. For genes, the thresholds of log2FC>1 and *P*adj<0.01 were used for upregulated genes (n=486), and the thresholds for log2FC<-0.01 and *P*adj<0.01 were used for downregulated genes (n=723), respectively. For retrotransposons, the thresholds of log2FC>3.5 and *P*adj<0.01 were used for upregulated retrotransposons (n=3,005), and the thresholds of log2FC<-3.5 and *P*adj<0.01 were used for downregulated retrotransposons (n=1,152), respectively. Upregulated and downregulated genes/retrotransposons are listed in **Supplementary Table 5**.

#### Gene Ontology and GSEA analyses

GO analyses for dysregulated genes and dysregulated SE-associated genes were performed using Metascape v3.5^99^. GO analyses for dysregulated typical enhancers and dysregulated RT-enhancers were performed using GREAT v4.0.4^100^. GSEA for dysregulated genes was performed using ClusterProfiler v4.6.2 with the Hallmark gene sets from MSigDB (Broad Institute)^101^.

#### Retrotransposon-derived transcript assembly

Retrotransposon-derived transcripts were assembled using StringTie2 v2.1.2 with default parameters^102^.

### ChIP-seq analyses

#### Mapping and visualization

Single-end sequencing reads were aligned to GRCh38/hg38 using ENCODE ChIP-seq pipeline v2.2.1 (https://github.com/ENCODE-DCC/chip-seq-pipeline2). Signal tracks with fold enrichment over input were generated using MACS2 v2.2.7.1 for visualization in genome browser^103^.

#### Peak calling and quality controls

Narrow peaks (H3K27ac and H3K4me3) were called using MACS2 v2.2.7.1 with parameters –g 2.7e9 –SPMR –keep-dup all and –q 0.01. High-confidence peaks were retained by further filtering peaks with (1) -log*P* > 5 and -log*Q* > 2, (2) input-subtracted RPM > 1 and (3) at least 2-fold enrichment of RPM in ChIP compared to input libraries (RPM_ChIP_ ≥ 2 × RPM_input_) as per ENCODE ChIP-seq pipeline (https://github.com/ENCODE-DCC/chip-seq-pipeline^2^). Broad peaks (H3K9me3 and H3K27me3) were called using epic2 v0.0.48^104^. For H3K9me3, the parameters –k –cs GRCh38.chrom.sizes –g 1 bin 5000 were used. For H3K27me3, the parameters –k –cs GRCh38.chrom.sizes –g 1 bin 2000 were used. For quality controls on ChIP-seq narrow peak datasets, NSC and RSC were calculated using phantompeakqualtools v1.2.2, and FRiP was calculated by dividing the number of reads falling into peak regions by the total number of mapped reads^105^.

#### Identification of CREs and dysregulated CREs

1-kb flanking region of H3K27ac and H3K4me3 summit peaks were used to identify putative CREs. Only experimental data are used to define putative CREs. Briefly, active putative enhancers were defined as regions enriched with H3K27ac peaks not overlapping with H3K4me3 peaks or GENCODE v39 TSS regions (±1kb flanking of GENCODE v39 TSS). Active putative promoters were defined as regions overlapped with both H3K27ac and H3K4me3 peaks. For samples that do not have H3K4me3 data, we defined active promoters as GENCODE v39 TSS regions that overlapped with H3K27ac, and active enhancers as H3K27ac peaks that do not overlap with active putative promoters. To generate a non-redundant union set of active enhancers or active promoters in HCC or TAN, the active enhancers or active promoters defined in each sample that are within 500 bp were first merged within sample, and once again across samples of the same condition. To generate a master list of active enhancers or active promoters across all samples, the non-redundant union set of active enhancers or promoters in HCC and TAN that are within 500 bp were merged. H3K27ac signals of all active enhancers (n=62,288) or active promoters (n=29,467) were subsequently subjected to k-means clustering using ComplexHeatmap v2.14.0^106^.

Dysregulated typical enhancers in HCC were defined using DESeq2 v1.36.0 with the thresholds log2FC > 2.25 and *P*adj<0.01 for activated typical enhancers (n=3,367), and log2FC < -1.5 and *P*adj<0.01 for inactivated typical enhancers (n=2,924), respectively. Similarly, dysregulated promoters in HCC were defined using DESeq2 v1.36.0 with the thresholds of log2FC>2 and *P*adj<0.01 for activated promoters (n=1,172), and log2FC<-2 and *P*adj < 0.01 for inactivated promoters (n=727), respectively. Dysregulated typical enhancers and promoters in HCC are listed in **Supplementary Table 6.**

#### Identification of SEs and dysregulated SEs

SEs were identified using the ROSE algorithm with default parameters. All SEs across samples (n=3,245) were generated using similar strategies as described above for differential analyses. Dysregulated SEs were defined using DESeq2 v1.36.0 with the thresholds of log2FC>1.5 and *P*adj<0.01 for activated SEs (n=234), and log2FC<-1.5 and *P*adj < 0.01 for inactivated SEs (n=86), respectively. Dysregulated SEs are listed in **Supplementary Table 6**.

#### Identification of dysregulated retrotransposon-derived CREs

Analyses were restricted to RepeatMasker-annotated retrotransposons (LTRs, LINEs and SINEs) that are at least 100 bp long and mapped to chr1-22 and chrX (n=3,195,734). Dysregulated RT-typical enhancers were defined as retrotransposons with at least 30% of its length overlapping with the dysregulated enhancers, and at least 15% of the dysregulated enhancers also overlapping with the retrotransposons, respectively (activated RT-typical enhancers, n=2,333; inactivated RT-typical enhancers, n=834). Dysregulated RT-promoters were similarly defined (activated RT-promoters, n=1,166; inactivated RT-promoters, n=195). Dysregulated RT-typical enhancers and RT-promoters are listed in **Supplementary Table 7**.

Over-represented subfamilies from activated RT-typical enhancers and activated RT-promoters were identified using the thresholds of observed/expected ratio ≥4, *Q* value (FDR-corrected one-tailed hypergeometric test) <0.01 and had at least 30 copies in the genome. The observed/expected ratio was calculated using the formula (n/n_all_)/(N/N_all_), where n is the number of a subfamily in activated RT-typical enhancers or activated RT-promoters, n_all_ is the total number of activated RT-typical enhancers or activated RT-promoters, N is the number of a subfamily in the genome and N_all_ is the total number of retrotransposons in the genome.

#### Motif analyses

Motifs in activated and inactivated typical enhancers were analyzed with HOMER v4.1.1^107^ findMotifsGenome.pl program using all non-redundant enhancers in HCC and TAN tissues (n=46,500) as background. Similar analyses were performed for activated RT-typical enhancers and activated RT-promoters using all RT-typical enhancers (n=24,596) and RT-promoters (n=10,261), respectively.

### Hi-C data analyses

HepG2 Hi-C data were downloaded from 4DN^108^. Significant chromatin interactions were called by FithiC2 v2.0.8 at 1-kb resolution with the threshold *Q*<0.05^109^.

### TCGA data analyses

#### Data download

TCGA Pan-Cancer cohort gene-level raw coverage count matrices, bigWig files and clinical metadata were downloaded using Recount3 v1.6.0^110^. TCGA-LIHC HM450 array data were downloaded from UCSC Xena^111^. HM450 probe annotation was downloaded from https://zwdzwd.github.io/InfiniumAnnotation. TCGA-LIHC GPC3 copy-number and mutation data were downloaded from cBioPortal^112–114^. TCGA-LIHC survival data were downloaded from TCGA PanCanAtlas Publications page at https://gdc.cancer.gov/about-data/publications/pancanatlas.

#### WGCNA of TCGA-LIHC cohort

WGCNA v1.71 was used to construct gene expression network in the TCGA-LIHC cohort^115^. Briefly, we filtered for genes that have at least 10 read counts in 90% of the TAN or HCC samples and excluded outlier samples. This resulted in a total of 26,108 genes across 408 samples. We then constructed the network using the blockwiseModules function using the parameters -power = 14, mergeCutHeight = 0.1 and deepSplit = 4, generating 84 modules. We calculated the correlation of each module to TAN and HCC and found three modules with high correlation to either TAN or HCC. Genes associated with the three modules were visualized using Gephi v0.10.1^116^.

#### DNA methylation level of GPC3 intragenic CGI and its relation to GPC3 expression

β-values for probes falling within the *GPC3* intragenic CGI (cg11442732, cg02442693 and cg09760728) were extracted using the HM450 probe annotation and correlated with *GPC3* expression in matched HCC tissues.

#### HERVE-int expression in the TCGA-PanCancer cohort

Raw reads falling on HERVE-int was quantified from TCGA Pan-Cancer bigWig files using Megadepth v1.8.0 and subsequently converted to TPM for analyses^117^.

#### Kapan-Meier survival analyses

Survival analyses for HERVE-int and THE1D were restricted to HCC patients with survival data. DESeq2-normalized counts of HERVE-int and THE1D were used for the analyses. For HERVE-int, patients were stratified into high-expression (top 25% percentile) and low-expression (bottom 25% percentile) groups. For THE1D, patients were stratified into high-expression (top 30% percentile), intermediate-expression (30%<x<70% percentile) and low-expression (bottom 30% percentile). Kaplan-Meier curves were generated using survminer v0.4.9 (https://github.com/kassambara/survminer) and survival v3.5.7 (https://github.com/therneau/survival). Patient stratification results are presented in Supplementary Table 8.

#### Multivariate cox regression analyses

Patients of the HERVE-int-High and HERVE-int-Low groups were binary categorized in the following clinicopathological features: alcohol consumption (no vs yes), HBV/HCV infection (no vs yes), NAFLD (no vs yes), tumor stage (I/II vs III/IV) and tumor grade (G1/2 vs G3/G4). Patients without the corresponding clinical annotation were assigned NA. Multivariate cox regression analyses were performed using survminer v0.4.9 and survival v3.5.7.

#### TMB analyses

TMB for HERVE-int-High and HERVE-int-Low patients was calculated with maftools v2.14.0 using the parameters: captureSize=35.8 and logScale=TRUE^118^.

### Publicly available data

All publicly available datasets used in this study are listed in **Supplementary Table 10.**

## Data availability

Epigenomic and transcriptomic datasets for the HCC and TAN samples were deposited to EGA under the accession ID EGA50000000035. H3K27ac and H3K4me3 ChIP-seq datasets for HepG2 and PLC were deposited to GEO under the accession code GSE276132. RNA-seq datasets for the CRISPRi experiments for HERVE-int and THE1D were deposited to GEO under the accession code GSE276133. All relevant source data are provided.

