## Supplementary materials for "Epigenomic analysis of hepatocellular carcinoma reveals aberrant *cis*-regulatory changes and dysregulated retrotransposons with prognostic potentials"

**This PDF file includes**:

Extended Data Figures 1-10

Supplementary Table 1

Legends for Supplementary Tables 1-10

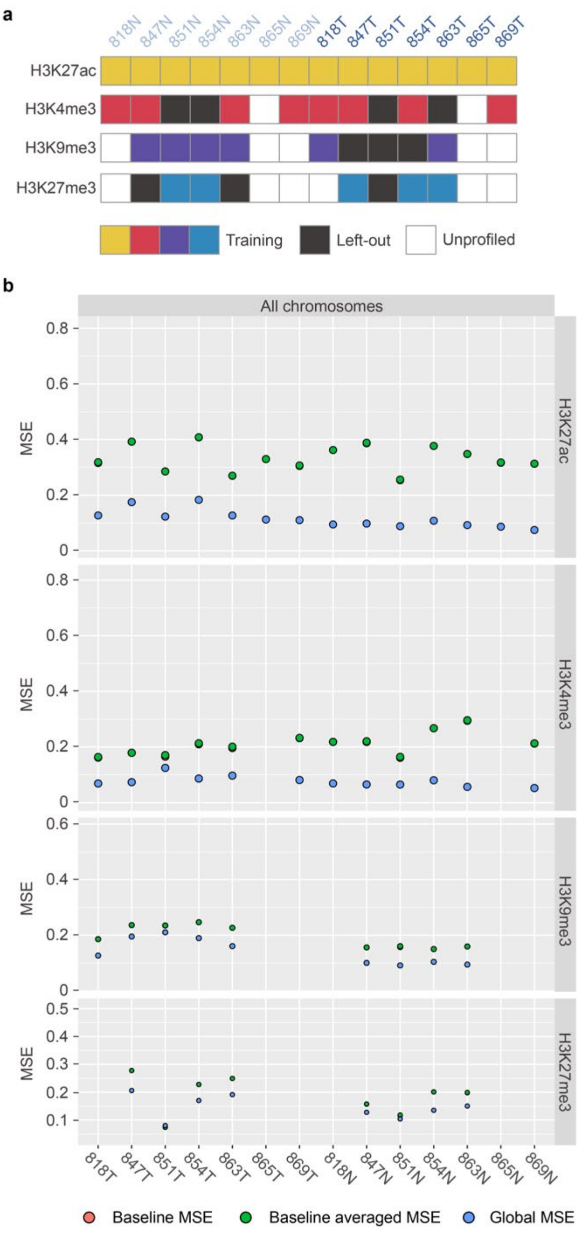


**Extended Data Fig.1. Quality controls of Avocado-imputed ChIP-seq signals. a**) Schematic of model training strategy. Models were trained on a per chromosome basis and 10 out of the 43 samples were left-out to assess the per-sample global mean-square error (MSE) of the imputed signal. **b**) Per-sample global MSE compared to per-sample baseline MSE and baseline MSE averaged across all other samples.

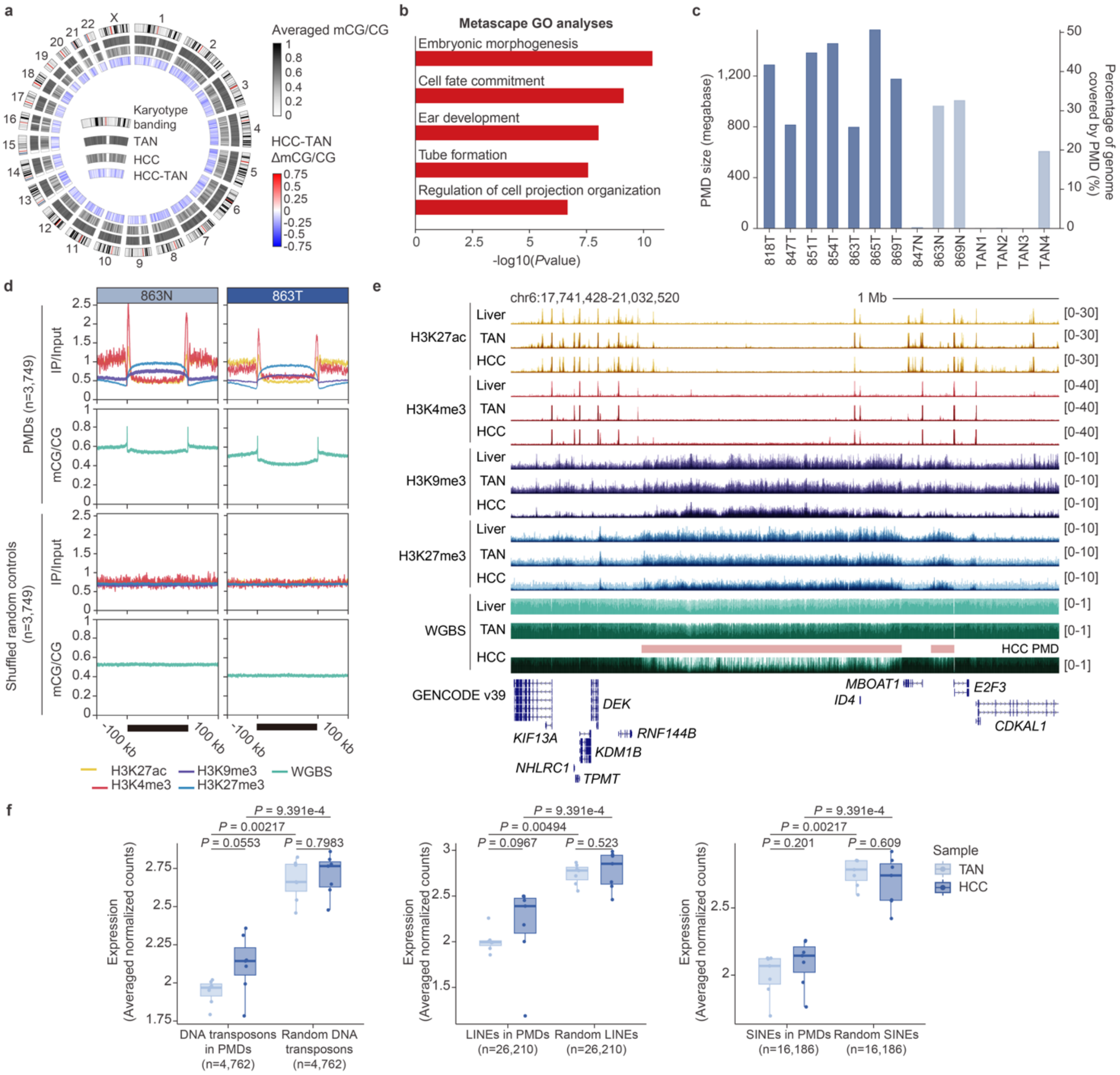


**Extended Data Fig.2. Characterization of DNA methylation alternations in HCC. a)** Circos plot showing the DNA methylation landscape of HCC and TAN. DNA methylation (mCG/CG) was calculated per megabase window and averaged across all HCC or TAN samples. The averaged DNA methylation difference (HCC-TAN ΔmCG/CG) between HCC and TAN samples is shown as heatmap. **b**) Gene ontology (GO) analyses of genes with hypermethylated promoters. *P* values (one-tailed hypergeometric test) for the top five enriched GO terms are shown. **c**) Barplots showing the PMD size and percentage of genome covered by PMD in each HCC or TAN sample. **d**) Aggregation plots of DNA methylation (mCG/CG) and histone mark signals (fold enrichment over input) in defined PMDs in patient 863. 100-kb flanking regions of defined PMDs are also shown. **e**) Genome browser screenshot showing two PMDs that are enriched with repressive histone marks, exhibit reduced DNA methylation and are depleted of active histone marks. PMDs are annotated as pink bars above the HCC WGBS track. **f)** Boxplots comparing the expression (averaged DESeq2-normalized counts) of DNA transposons (left), LINEs (middle) and SINEs (right) between HCC and TAN that are localized within HCC PMDs. *P* values (two-tailed Wilcoxon test) are shown. The center and bounds of boxes indicate the median and quartile of all data points, respectively. The minima and maxima of whiskers indicate quartile $1 - 1.5 \times$ the interquartile range and quartile $3 + 1.5 \times$ the interquartile range, respectively.

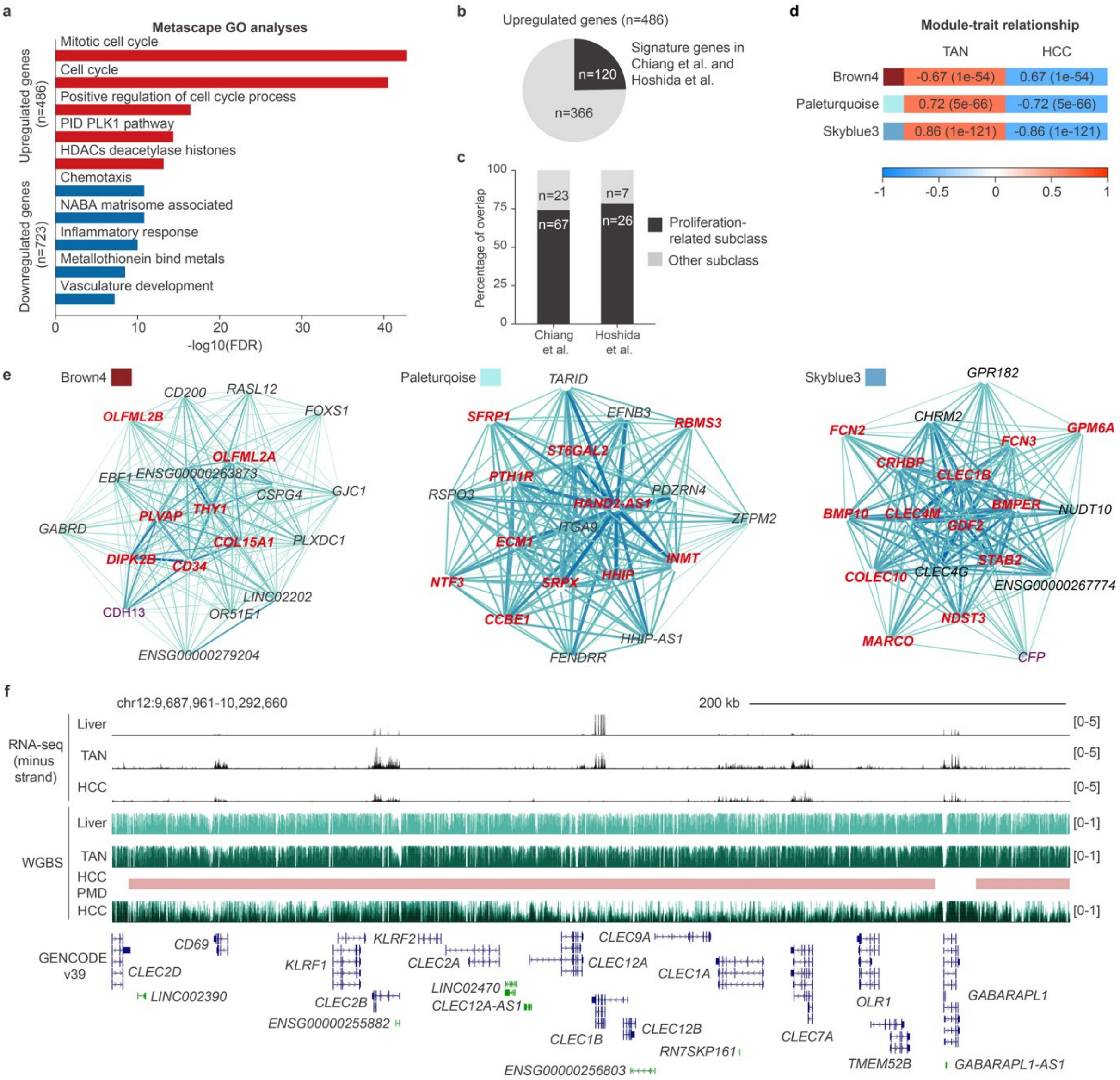


**Extended Data Fig.3. Characterization of transcriptomic dysregulations in HCC. a**) GO analyses for dysregulated genes in HCC. *P* values (one-tailed hypergeometric test) for the top 5 GO terms are shown for both the upregulated and downregulated genes, respectively. **b**) Pie chart showing the number of upregulated genes defined as signature genes under the molecular subclass classifications by Chiang *et al*. and Hoshida *et al*. **c**) Barplots showing the number of signature genes in **b** associated with cell proliferation pathways under the molecular subclass classifications by Chiang *et al*. and Hoshida *et al*. Note that 3 upregulated genes were classified as signature genes by both Chiang *et al*. and Hoshida *et al*. **d**, WGCNA modules significantly associated with HCC (Brown4) or TAN (Paleturqoise and Skyblue3). **e**, Network graphs of modules significantly associated with HCC (Brown 4, left) or TAN (Paleturqoise, middle and Skyblue3, right). Dysregulated genes defined in our HCC samples are highlighted in red within each module. In the network, node represents a gene, and each edge represents the co-expression relationship between two genes. **f**, Genome browser screenshot showing the transcriptional downregulation of *CLEC1B* coincides with loss of DNA methylation in PMD. PMDs in HCC are annotated as pink bars above the HCC WGBS track. Stranded total RNA-seq tracks are displayed as TPM, and WGBS tracks are displayed as mCG/CG. Liver RNA-seq and WGBS tracks are from Roadmap Epigenomics Project and shown as single replicates. Other tracks are shown as composite signals by overlaying tracks across samples.

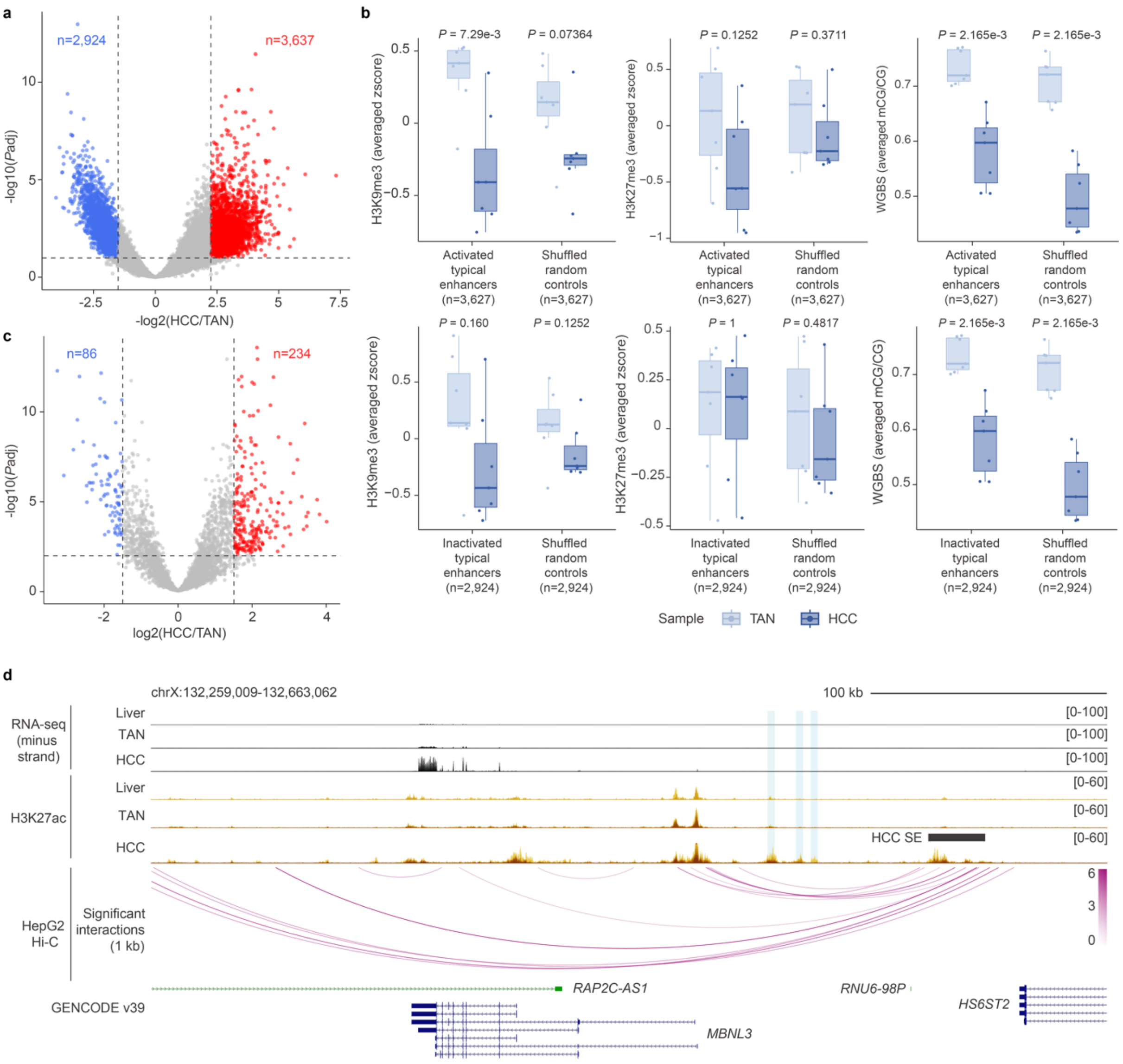


**Extended Data Fig. 4. Characterization of dysregulated putative enhancers and super-enhancers (SEs) in HCC. a**) Volcano plot showing dysregulated typical enhancers in HCC. For each typical enhancer, the -log10-transformed, two-tailed adjusted *P* values (*P*adj) is plotted against the log2-transformed fold change (HCC/TAN). Activated **(***P*adj < 0.01 and log2FC > 2.25; n=3,637**)** and inactivated typical enhancers (*P*adj < 0.01 and log2FC < −1.5; n=2,924) are labeled in red and blue, respectively. Dashed lines represent the indicated thresholds. **b**) Boxplots comparing the averaged H3K9me3 (left) and H3K27me3 (middle) signals (z-score transformed -log(*P*value) signal), and averaged DNA methylation (mCG/CG) of dysregulated typical enhancers between HCC and TAN. Each dot represents a sample. *P* values (two-tailed Wilcoxon test) are shown. **c**) Volcano plot showing dysregulated SEs in HCC.

For each SE, the -log10-transformed two-tailed adjusted *P* values (*P*adj) is plotted against the log2-transformed fold change (HCC/TAN). Activated (*P*adj < 0.01 and log2FC > 1.5) and inactivated SEs (*P*adj < 0.01 and log2FC < −1.5) are labeled in red and blue, respectively. Dashed lines represent the indicated thresholds. **d**) Genome browser screenshot showing an activated SE and several activated typical enhancers in HCC at the *MBNL3* locus. The activated SE is annotated as gray block above the HCC H3K27ac track, and the activated typical enhancers are highlighted by cyan shadings. Significant interactions defined from HepG2 Hi-C are shown as arcs and the color denotes significance (-log(*Q*value)). Stranded RNA-seq tracks are displayed as TPM, and H3K27ac ChIP-seq tracks are displayed as fold enrichment over input. Liver RNA-seq track is from Roadmap Epigenomics Project and is shown as single replicate. Other tracks are shown as composite signals by overlaying tracks across samples.

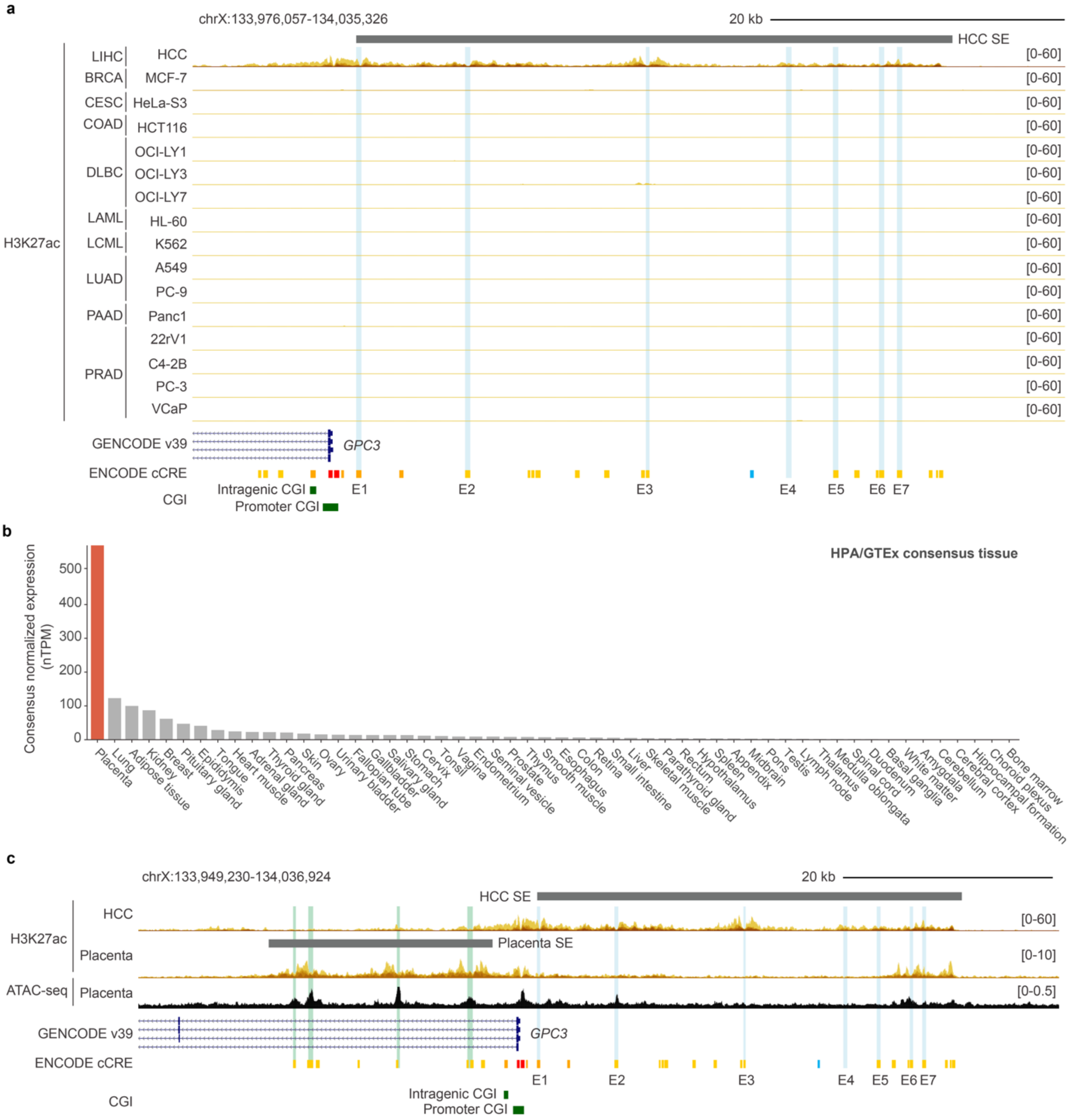


**Extended Data Fig. 5. Cell type-differential SE for *GPC3* in HCC and placenta. a)** Genome browser screenshot showing the absence of H3K27ac signals at the *GPC3*-associated SE in other cancer cell lines representing different cancer types. Cancer type abbreviations follow TCGA definitions. The constituent enhancers (E1-E7) of the *GPC3* SE are highlighted by cyan shadings. **b**) *GPC3* expression in normalized TPM across a panel of normal tissues from Human Protein Atlas (https://www.proteinatlas.org). **c**) Genome browser screenshot showing cell type-differential SEs for *GPC3* transcriptional regulation in HCC and placenta. SE definitions are shown as gray bars above their respective H3K27ac tracks. The constituent enhancers of the *GPC3* SE in HCC and that in placenta are highlighted by cyan and green shadings, respectively. Placenta ATAC-seq data were from Gao, *et al*..

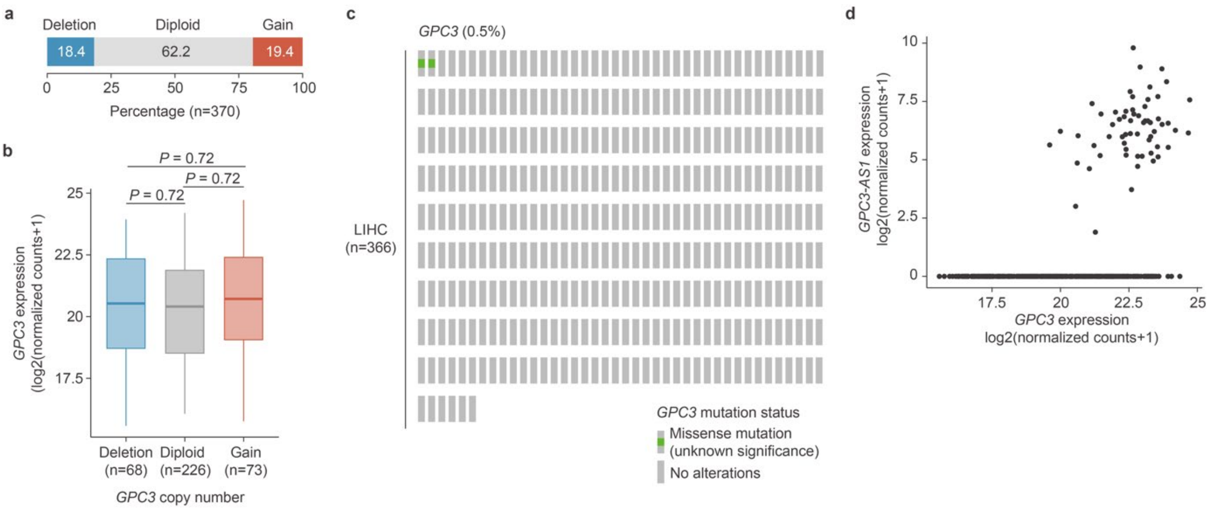


**Extended Data Fig.6.** ***GPC3* upregulation in HCC cannot be explained by copy number alternations, mutations nor overexpression of *GPC3-AS1*. a**) Copy number status of *GPC3* in the TCGA-LIHC cohort. **b**) Boxplots comparing the expression of *GPC3* (log2-transformed, DESeq2-normalized counts) among HCC patients with different copy number status of *GPC3* in the TCGA-LIHC cohort. **c**) Mutation status of *GPC3* in the TCGA-LIHC cohort. **d**) Scatterplot showing the expression correlation between *GPC3* and *GPC3-AS1* (log2-transformed, DESeq2-normalized counts) in the TCGA-LIHC cohort. Each dot represents a patient.

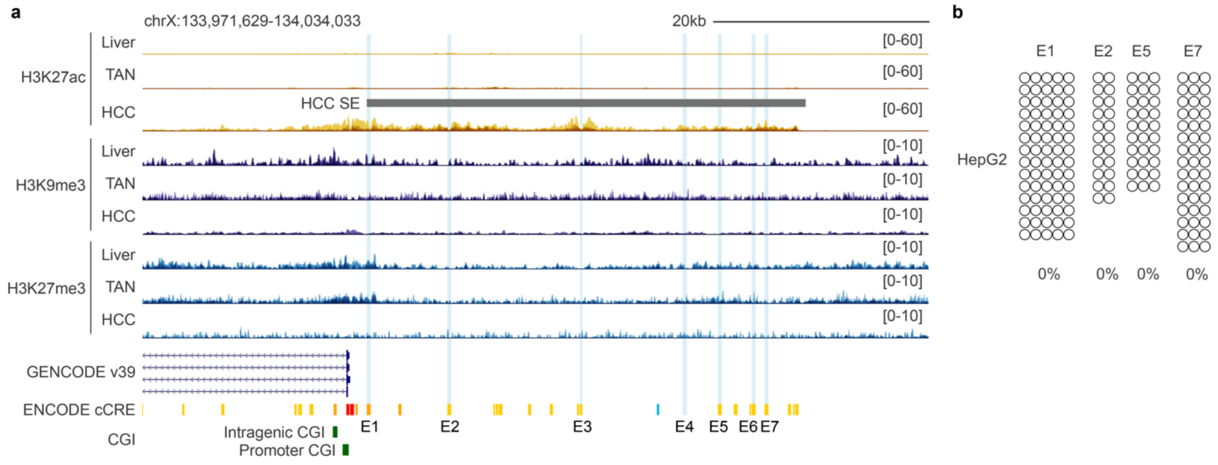


**Extended Data Fig. 7. Reactivation of *GPC3*-associated SE in HCC is associated with loss of DNA methylation but not H3K9me3 nor H3K27me3. a**) Genome browser screenshot showing the subtle changes in H3K9me3 and H3K27me3 at the *GPC3*-associated SE. The SE is annotated as gray bar above the H3K27ac track. The constituent enhancers (E1-E7) of the *GPC3* SE are highlighted by cyan shadings. ChIP-seq signals are displayed as fold enrichment over input. All tracks are shown as composite signals by overlaying tracks across samples. **b**) Bisulfite amplicon sequencing results for constituent enhancers of *GPC3* SE that harbour at least two CpGs (E1, E2, E5 and E7). Hollow circle indicates unmethylated CpG.

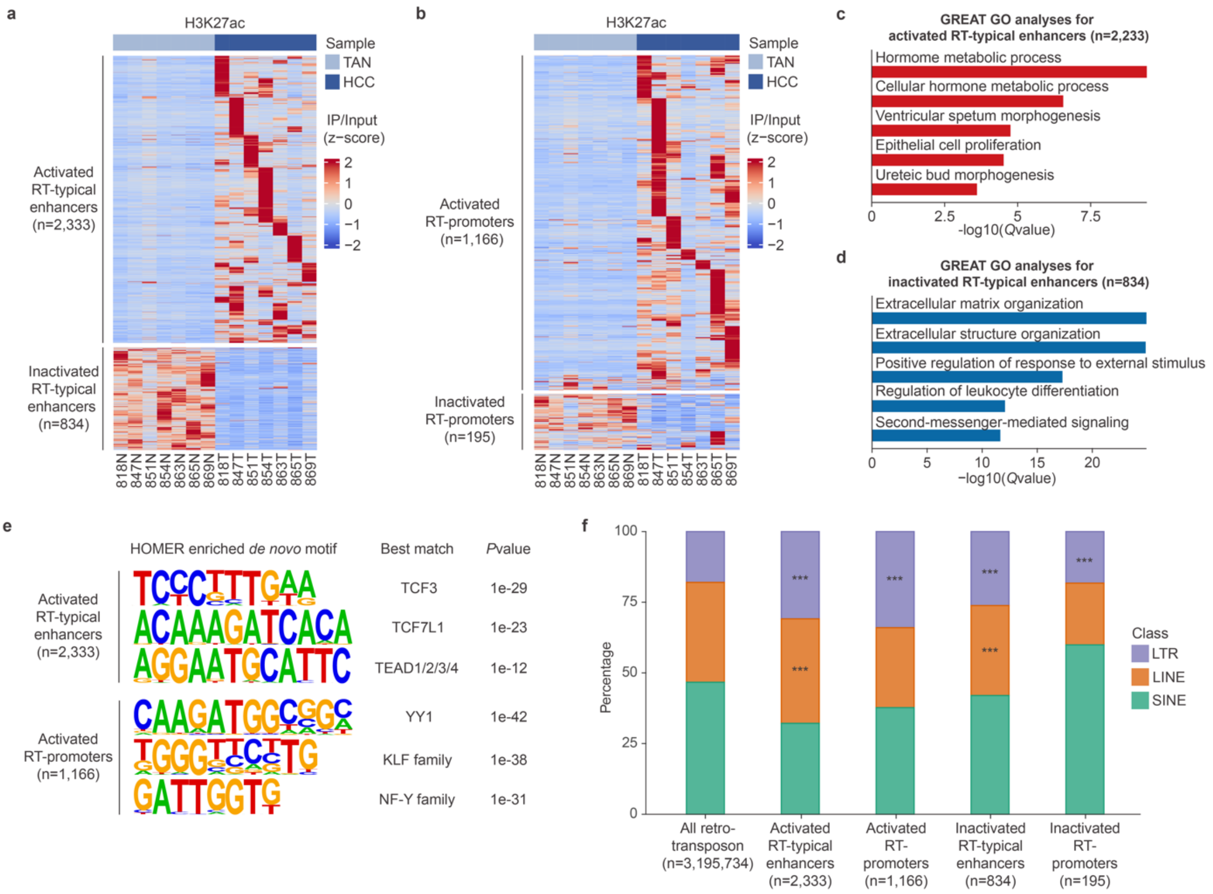


**Extended Data Fig.8. Epigenetic dysregulation of retrotransposon-derived CREs in HCC. a-b)** Heatmap of H3K27ac signals (z-score transformed fold enrichment over input) of dysregulated RT-typical enhancers (**a**) and RT-promoters (**b**) in HCC. Each row represents a dysregulated element, and each column represents a sample. **c**) Activated RT-typical enhancers were associated with genes related to metabolism and early development. **d**) Activated RT-promoters were associated with extracellular matrix organization. **c-d**) *Q* values (Binomial test) for the top 5 enriched GO terms are shown. **e**) Motif analyses revealing enrichment of binding motifs for developmental TFs among activated RT-typical enhancers, and binding motifs for ubiquitously expressed TFs among activated RT-promoters. **f**) Barplots comparing the percentage of the three classes of retrotransposon among the dysregulated retrotransposon-derived CREs. Activated RT-typical enhancers: LTR ($P$ = 2.41e-131), LINE ($P$ = 1.12e-41); activated RT-promoters: LTR ($P$ = 2.442e-87); inactivated RT-typical enhancers: LTR ($P$ = 4.760e-51), LINE ($P$ = 1.47e-4); inactivated RT-promoters: LTR ($P$ = 2.487e-5). $P$ values (one-tailed hypergeometric test corrected by the Benjamini-Hochberg approach) are shown.

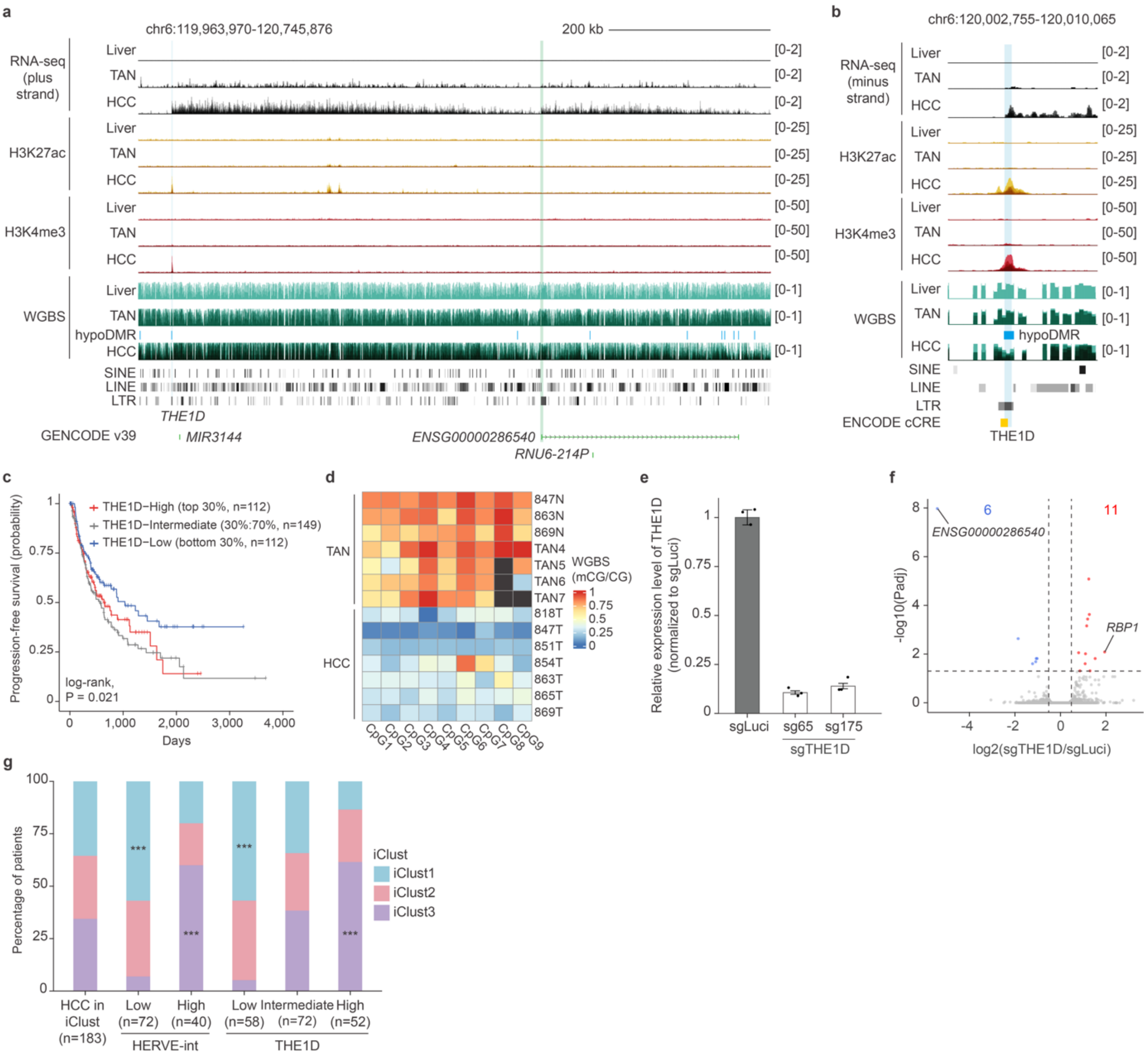


**Extended Data Fig.9. Epigenetic dysregulation of THE1D-driven transcripts in HCC.** **a-b**) Genome browser screenshot showing THE1D serving as the alternative promoter for the pseudogene *ENSG00000286540* in HCC (**a**) and a zoom-in screenshot of the THE1D element gaining active promoter histone marks (**b**). THE1D is highlighted by cyan shading and the *ENSG00000286540* pseudogene promoter is highlighted by green shading, respectively. HypoDMR in HCC is annotated as blue bar above the HCC WGBS track. Stranded total RNA-seq tracks are displayed as TPM, ChIP-seq tracks are displayed as fold enrichment over input, and WGBS tracks are displayed as mCG/CG. Liver RNA-seq and WGBS tracks are from Roadmap Epigenomics Project and shown as single replicates**.** Other tracks are shown as composite signals by overlaying tracks across samples. **c**) Kaplan-Meier survival curve for

THE1D expression among HCC patients in the TCGA-LIHC cohort. Based on the THE1D expression in DESeq2-normalized counts, patients were stratified into low (bottom 30%), intermediate (30%<x<70%) and high (top 30%), groups, respectively. *P* value (log-rank test) is shown. **d**) Heatmap comparing the DNA methylation (mCG/CG) of THE1D between HCC and TAN. Each row represents a sample, and each column represents a CpG within THE1D. Dark gray cells indicate CpGs with insufficient read coverage (< 5). **e**) Boxplots showing the relative expression of THE1D measured by RT-qPCR in HepG2 cells transduced with THE1D-targeting sgRNAs (sgTHE1D) or luciferase-targeting sgRNA control (sgLuci). Each dot represents a technical replicate and error bars denote standard deviations with the centers indicating the means of three technical replicates. *GAPDH* is used as internal control and relative THE1D expression is normalized to HepG2 cells transduced with sgLuci. **f**) Volcano plot showing dysregulated genes in HepG2 cells depleted of THE1D. For each gene, the -log10-transformed, two-tailed adjusted *P*values (*P*adj) is plotted against the log2-transformed change (sgTHE1D/sgLuci). Upregulated (*P*adj < 0.05 and log2(Fold change (sgTHE1D/sgLuci) > 0.5) and downregulated (*P*adj < 0.05 and log2(Fold change (sgTHE1D/sgLuci) < -0.5) are labeled in red and blue, respectively. Dashed lines represent the indicated thresholds. Selected dysregulated genes are labeled. **g**) Barplots comparing the percentage of HERVE-int-Low/-High and THE1D-Low/-Intermediate/-High patients annotated with the iCluster1-3 definitions from TCGA. HERVE-int-Low: iClust1 (*P* = 7.148e-6); HERVE-int-High: iClust3 (*P* = 4.915e-4); THE1D-Low: iClust1 (*P* = 2.045e-4). THE1D-High (*P* = 1.675e-5). **P* < 0.05 and ****P* < 0.001, one-tailed hypergeometric test corrected by the Benjamini-Hochberg approach.

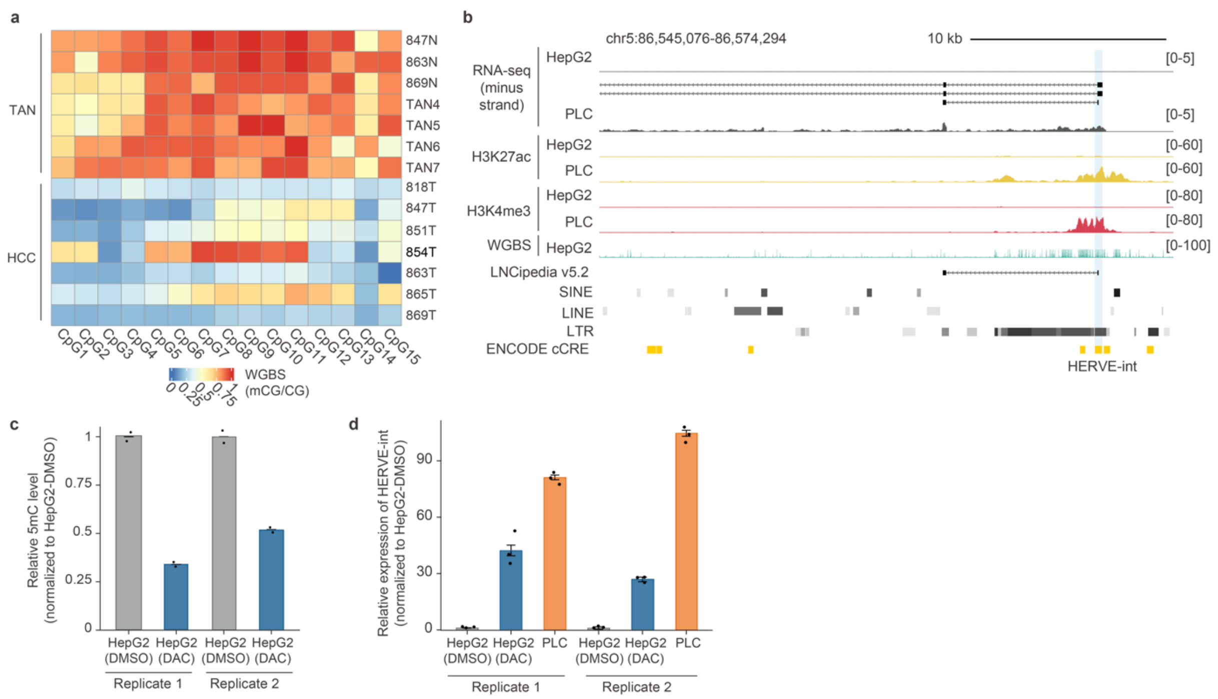


**Extended Data Fig. 10. DNA hypomethylation is sufficient for HERVE-int de-repression in HCC. a**) Heatmap comparing the DNA methylation (mCG/CG) of HERVE-int between HCC and TAN. Each row represents a sample, and each column represents a CpG within HERVE-int. **b**) Genome browser screenshot showing the differential expression of HERVE-int in PLC cells. StringTie-assembled transcripts are shown above the HCC RNA-seq track. HERVE-int is highlighted by cyan shading. All tracks for HepG2 and PLC are shown as single replicate. **c**) Boxplots showing the relative DNA methylation level in HepG2 cells treated with dimethyl sulphoxide (DMSO) or 5-aza-2'-deoxycytidine (DAC) in two independent experiments. Each dot represents a technical replicate. **d)** Boxplots showing the relative expression of HERVE-int measured by RT-qPCR in HepG2 cells treated with DAC or DMSO in two independent experiments. Each dot represents a technical replicate and error bars denote standard deviations with the centers indicating the means of three technical replicates. *GAPDH* is used as internal control and relative HERVE-int expression is normalized to HepG2 cells treated with DMSO.

**Legends for Supplementary Tables**

**Supplementary Table 1. Clinical metadata of enrolled HCC patients**

| Patient | Etiology | Age | Gender | Cirrhosis | Steatosis | Vascular invasion | Smoker | Drinker | Differentiation | Number of tumors | Size (cm) |
|---|---|---|---|---|---|---|---|---|---|---|---|
| 818 | HBV | 57 | M | Yes | Yes | No | No | No | Moderate | 1 | 3.5x3.5x2 |
| 847 | HBV | 66 | F | No | No | Yes | Ex | Ex | Moderate | Multiple | up to 7 |
| 851 | HBV | 58 | M | Yes | Yes | No | No | No | Well | 1 | 2.5 |
| 854 | HBV | 55 | M | Yes | Yes | Yes | Ex | No | Moderate | 1 | 4.5x2.5x2.5 |
| 863 | HBV | 51 | M | Yes | No | Yes | No | No | Moderate | 2 | 2.5, 1.5 |
| 865 | HBV | 71 | M | Yes | No | No | No | No | Well | 1 | 6x5x5.5 |
| 869 | HBV | 54 | M | Yes | Yes | Yes | Current | No | Moderate | 3 | 2.5x2.2x2, 1.6, 0.8 |

**Supplementary Table 2.** **Quality metrices for WGBS and ChIP-seq datasets.**

**Supplementary Table 3.** **List of oligos.**

**Supplementary Table 4.** **List of DMRs and PMDs in HCC.**

**Supplementary Table 5.** **List of dysregulated genes and retrotransposons in HCC.**

**Supplementary Table 6.** **List of dysregulated promoters, typical enhancers and SEs in HCC.**

**Supplementary Table 7.** **List of dysregulated RT-promoters and RT-typical enhancers in HCC.**

**Supplementary Table 8.** **HERVE-int and THE1D expressions in the TCGA-LIHC cohort.**

**Supplementary Table 9.** **List of dysregulated genes in HERVE-int-depleted PLC cells and THE1D-depleted HepG2 cells.**

**Supplementary Table 10.** **List of public data.**

Supplementary Tables 2-10 are provided in separate excel spreadsheets.
